# MAIT cells drive chronic inflammation in a genetically diverse murine model of spontaneous colitis

**DOI:** 10.1101/2023.11.29.569225

**Authors:** Liyen Loh, David Orlicky, Andrea Spengler, Cassandra Levens, Sofia Celli, Joanne Domenico, Jared Klarquist, Joseph Onyiah, Jennifer Matsuda, Kristine Kuhn, Laurent Gapin

**Affiliations:** Department of Immunology and Microbiology, University of Colorado School of Medicine, Aurora, CO, USA; Department of Pathology, University of Colorado School of Medicine, Aurora, CO, USA; Division of Rheumatology, Department of Medicine, University of Colorado School of Medicine, Aurora, CO, USA; Division of Gastroenterology, Department of Medicine, University of Colorado School of Medicine, Aurora, CO, USA; Mouse Genetics Core, National Jewish Health, Denver, CO, USA

**Author notes:** Corresponding Authors, Laurent Gapin, Liyen Loh. **Grant Support**: National Institutes of Allergy and Infectious Diseases R21 AI156003 (L.G), R21 AI163454 (L.G), University of Colorado Gastrointestinal and Liver Innate Immune Program pilot grant (L.L), University of Colorado RNA Bioscience Initiative Award (L.L). **Abbreviations:** interleukin (IL), Mucosal-Associated Invariant T (MAIT), single cell RNA sequencing (scRNAseq), collaborative cross (CC), innate lymphoid cells (ILC3), unconventional T (T_UNC_), T cell receptor (TCR), major histocompatibility complex (MHC), MHC-class I-related (MR1), inflammatory bowel disease (IBD), quantitative trait locus (QTL), invariant natural killer T (iNKT), promyelocytic leukemia zinc finger (PLZF), retinoic acid related orphan receptor-gamma (ROR-γt), hematoxylin and eosin (H&E), polymorphonuclear (PMN), lamina propria (LP), regulatory T (Treg), T helper (Th), conventional T (T_CONV_), genome-wide association studies (GWAS). **CReDiT Author Contributions**: Liyen Loh, PhD (Conceptualization: Equal; Formal analysis: Lead; Funding acquisition: Supporting; Investigation: Lead; Methodology: Lead; Writing – original draft: Equal; Writing – review & editing: Equal) David Orlicky, PhD (Formal analysis: Supporting; Writing – review & editing: Supporting) Andrea Spengler, BS (Formal analysis: Supporting; Investigation: Supporting; Writing – review & editing: Supporting) Cassandra Levens, (Investigation: Supporting, Writing – review & editing: Supporting) Sofia Celli, BA (Formal analysis: supporting, Methodology: Supporting; Writing – review & editing: Supporting) Joanne Domenico, BA (Investigation: Supporting; Writing – review & editing: Supporting) Jared Klarquist, PhD (Formal analysis: Supporting; Investigation: Supporting; Writing – review & editing: Supporting) Joseph Onyiah, MD, (Methodology: Supporting; Writing – review & editing: Supporting) Jennifer Matsuda, PhD (Investigation: Supporting; Methodology: Supporting; Writing – review & editing: Equal Supporting) Kristine Kuhn, MD, PhD (Methodology: Supporting; Funding acquisition: Supporting; Writing – review & editing: Supporting) Laurent Gapin, PhD (Conceptualization: Equal; Funding acquisition: Lead; Methodology: Supporting; Writing – original draft: Equal; Writing – review & editing: Equal). **Data transparency statement**: Data that support the findings of this study will be deposited in NCBI GEO with an accession code. R Scripts for analyses will be available on github.

**Keywords:** MAIT cells, colitis, inflammation, colon, IL-17, scRNAseq

## Abstract

**Background & aims:** Lymphocytes that produce IL-17 can confer protective immunity during infections by pathogens, yet their involvement in inflammatory diseases is a subject of debate. Although these cells may perpetuate inflammation, resulting in tissue damage, they are also capable of contributing directly or indirectly to tissue repair, thus necessitating more detailed investigation. Mucosal-Associated-Invariant-T (MAIT) cells are innate-like T cells, acquiring a type III phenotype in the thymus. Here, we dissected the role of MAIT cells *in vivo* using a spontaneous colitis model in a genetically diverse mouse strain.

**Methods:** Multiparameter spectral flow cytometry and scRNAseq were used to characterize MAIT and immune cell dynamics and transcriptomic signatures respectively, in the collaborative-cross strain, CC011/Unc and CC011/Unc-*Traj33^-/-^*.

**Results:** In contrast to many conventional mouse laboratory strains, the CC011 strain harbors a high baseline population of MAIT cells. We observed an age-related increase in colonic MAIT cells, Th17 cells, regulatory T cells, and neutrophils, which paralleled the development of spontaneous colitis. This progression manifested histological traits reminiscent of human IBD. The transcriptomic analysis of colonic MAIT cells from CC011 revealed an activation profile consistent with an inflammatory milieu, marked by an enhanced type-III response. Notably, IL-17A was abundantly secreted by MAIT cells in the colons of afflicted mice. Conversely, in the MAIT cell-deficient CC011-Traj33−/− mice, there was a notable absence of significant colonic histopathology. Furthermore, myeloperoxidase staining indicated a substantial decrease in colonic neutrophils.

**Conclusions:** Our findings suggest that MAIT cells play a pivotal role in modulating the severity of intestinal pathology, potentially orchestrating the inflammatory process by driving the accumulation of neutrophils within the colonic environment.

## Introduction

The IL-17 cytokine family, encompassing IL-17A through IL-17F, plays a crucial role in triggering inflammation in the face of pathogenic microbial threats, thereby conferring protection^1, 2^. Deficiencies in the IL-17 signaling pathway, due to loss-of-function mutations in associated genes, result in a predisposition to persistent or recurrent infections, both fungal and bacterial^3, 4^. On the other hand, levels of IL-17 are heightened in individuals with various inflammatory and some autoimmune conditions^5, 6^, where it is believed to exacerbate immunopathology, particularly when acting in concert with other pro-inflammatory cytokines such as IL-23 and IL-6^2^. Notably, the monoclonal IgG1 antibody Secukinumab, which inhibits IL-17, has been effectively employed in the treatment of psoriasis^7^ and rheumatoid arthritis^8^. Nonetheless, its application has led to unexpected and adverse outcomes in patients with active inflammatory bowel diseases, including Crohn’s disease^9^. The immune mechanisms underlying these paradoxical effects are not well defined.

IL-17 is secreted by a variety of cell types spanning both the innate and adaptive immune systems, including macrophages, Group 3 innate lymphoid cells (ILC3), lymphoid tissue-inducer cells, Th17 cells, and subsets of unconventional T (T_UNC_) cells that recognize non-peptide antigens^2^. These cells are unified by the expression of ROR-γt, the master transcription factor defining the IL-17-producing lineage, and they predominantly localize at barrier tissues. While Th17 cells are well-known contributors to inflammation in a host of autoimmune and inflammatory disorders^2, 10^, Mucosal-associated invariant T (MAIT) cells have recently emerged as significant players in IL-17-driven immunity, endowed with a versatile cytokine-producing capability acquired during thymic development^11–15^. Their prominence in human physiology is underscored by their abundance in human blood and tissues, notably within the gastrointestinal tract, where they can constitute up to 6% of T cells in the colon^16–19^, as well as their evolutionary conservation across species^20–22^.

The MAIT cell T cell receptor (TCR) is remarkably conserved among individuals, recognizing microbial-derived metabolite antigens presented by MR1, a non-classical and non-polymorphic MHC molecule. Activation of the MAIT TCR culminates in a pro-inflammatory response, implicating these cells in the pathology of autoimmune diseases such as IBD^11, 23, 24^. Patients with IBD demonstrate significant infiltration of MAIT cells into inflamed gut tissue where they display an inflammatory profile^25–27^. Although, the possible correlations between their frequencies and involvement towards colitis disease remains uncertain. Murine IBD models have yielded mixed findings regarding whether MAIT cells exacerbate^28^ or mitigate^29^ colitis. The advent of single-cell transcriptomics has shed light on the shared traits between MAIT cells and the Th17 lineage, revealing that human blood MAIT cells acquire a wound repair signature following TCR and cytokine activation. *In vivo* studies in barrier tissues like the skin have translated these transcriptional patterns into functional roles in tissue repair and wound healing^30, 31^. This emerging evidence hints at the dualistic nature of MAIT cells in inflammatory diseases, underscoring the need for more research into the nuanced and tissue-specific roles of MAIT cells in the pathogenesis of IBD.

The susceptibility to IBD has been traced to numerous genetic loci and is further complicated by environmental influences, such as the gut microbiota^32, 33^. Traditional *in vivo* models of colitis typically induce colitis using chemical agents or pathogens, which may not accurately replicate the progression of human disease due to the rapid onset and potential for fatal outcomes^34^. While there are models of spontaneous colitis, they usually involve single gene deletions, and IBD rarely results from monogenic defects^35^. The Collaborative Cross (CC) resource has provided construction of a genetically diverse mouse population, designed to enhance our understanding of human diseases^36^. Within the CC strains, extreme phenotypes can be identified, and the responsible genetic loci pinpointed through quantitative trait locus (QTL) mapping, thanks to the well-documented genetic makeup of these strains^36^. Notably, one particular CC strain, the CC011/Unc (CC011), has been observed to spontaneously develop colitis past 20 weeks of age in the absence of any known pathogens^37^. Although previous QTL mapping linked the colitis phenotype to four specific loci^37^, the immunological factors contributing to the disease have yet to be investigated.

In this study, we harnessed the CC011 mouse line to explore the involvement of MAIT cells in the spontaneous development of colitis. In a striking contrast to typical laboratory strains, where MAIT cells are seldom present, we found that young CC011 mice have a significantly higher prevalence of MAIT cells—about 15 times greater than the average seen in 50 other CC strains and the 8 original founder strains—in both thymic and peripheral tissues.

Aligning with the observations made by Rogala et al.^37^, we noted the development of the typical colitis phenotype within the CC011 mouse cohort, a condition that manifested with an age-dependent increase of MAIT cells, Th17 cells, and neutrophils in the colon. Utilizing single-cell RNA sequencing (scRNAseq), we identified that MAIT cells from the colons of afflicted mice displayed a transcriptome enriched with markers for Th17 lineage differentiation and tissue repair capabilities, in addition to activation signals triggered by the surrounding inflammatory context. Furthermore, these colonic MAIT cells were capable of producing IL-17A in affected animals *in vivo*.

The generation of a MAIT cell-deficient Traj33−/− strain on the CC011genetic background unveiled a milder colonic histopathology in older mice, alongside a noticeable decline in neutrophil presence within both the colon and extra-intestinal tissues. These results underscore the pathogenic involvement of MAIT cells in colitis pathogenesis and highlight their significance as mediators of intestinal inflammation.

## Results

### CC011 a MAIT cell-rich collaborative cross strain at steady-state

Although the CC mouse strains offer a genetically varied and defined platform for the study of human diseases, the distribution of MAIT cells within these strains has not been characterized. We analyzed the frequency of MAIT cells in the thymus of 6-9 week-old mice across the eight founder strains (A/J, C57BL/6J (B6), 129S1/SvImJ, NOD/SHiLtJ, NZO/HiltJ, CAST/EiJ, PWK/PhJ, and WSB/Eij) as well as 50 other CC strains. We employed MR1-5-OP-RU-APC tetramer staining alongside TCR-β detection to accurately identify MAIT cells (Figure 1A, Supplementary Figure 1). Among the 58 strains assessed, CC011 stood out, having approximately an 18-fold increase in MAIT cells compared to many of the strains (Figure 1A/B), with the exception of the CC030/GeniUnc strain, which exhibited the second-highest MAIT cell frequency (Figure1B, Supplementary Table 1).

**Figure 1:**
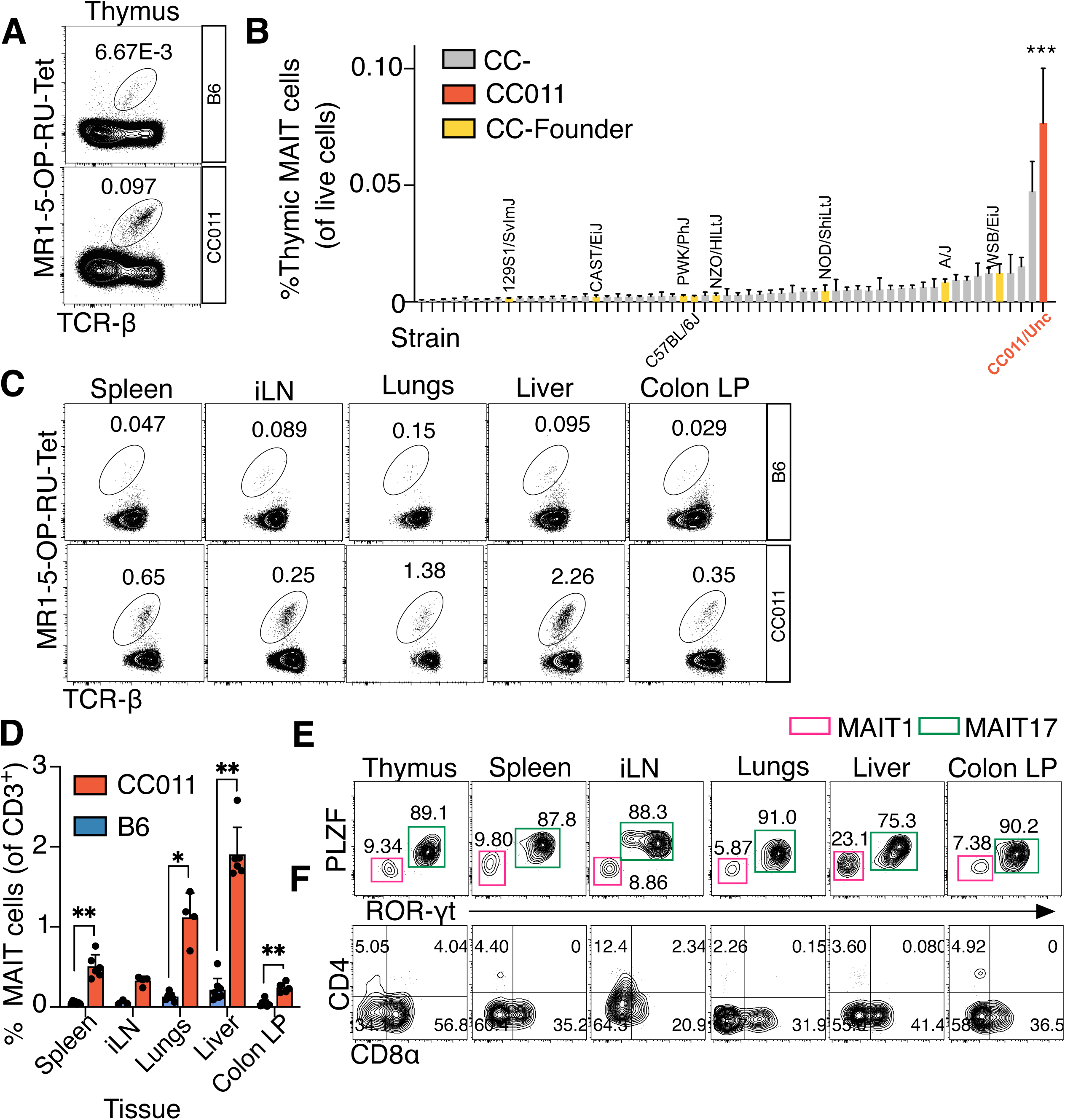
CC011 a MAIT cell-rich collaborative cross strain at steady-state. A. Representative contour flow cytometry plots of MAIT cells in CC011 and founder reference strain B6, shown is the % frequency of MAIT cells of total live cells in the thymus. B. Summarized frequencies of A (Mean, SD) across 50 CC strains and 8 founder strains, the table of strains is shown in Table S1. C. Representative contour flow cytometry plots of MAIT cells in the tissues of B6 and CC011 mice, shown is the % frequency of MAIT cells of CD3^+^ T cells. D. Summary of C, where circles represent individual mice from B6 (Blue) or CC011 (Red) strains. E. Intracellular staining for transcription factors and F. cell surface co-receptors in CC011 mice. Shown is the % frequency of the transcription factor or co-receptor of MAIT cells. (* P<0.05; **P<0.01; ***P<0.001; ****P<0.0001, One-way ANOVA, multiple comparisons to CC011 (B), n=2-6 mice/group; Mann-Whitney Test, (D) n= 3-8 mice/group all mice were age 6-9 weeks old).

For comparative purposes, we used the B6 strain as a control, given its status as one of the CC’s founding strains and its lack of spontaneous colitis development^37^. In both lymphoid and peripheral tissues, CC011 mice showed a pronounced prevalence of MAIT cells in contrast to B6 mice (Figure 1C/D). Notably, the greatest percentages of MAIT cells were detected in the lungs and liver, constituting approximately 1-1.5% of T cells (Figure 1D). In previous studies, an increase in tissue-resident MAIT cell frequencies in the absence of other unconventional T cell subsets, such as iNKT or γ8 T cells, has been noted, indicating a compensatory or competitive relationship^14, 30, 38^. In the thymus of the CC011 strain, we observed a notable decrease in the frequency of iNKT cells in comparison to the B6 mice, while the prevalence of γ8 T cells appeared equivalent between the two strains (Supplementary Figure 2A). This pattern was consistent in other tissues as well: iNKT cell frequencies were similarly diminished in both the spleen and liver of the CC011 mice (Supplementary Figure 2B). Conversely, γ8 T cells were found in greater numbers within the spleens of the CC011 strain relative to their B6 counterparts (Supplementary Figure 2C). This could suggest an interplay of niche competition among these cell populations.

In the CC011 strain, the distribution of the key transcription factors for T_UNC_ cells and IL-17-producing cells, promyelocytic leukemia zinc finger (PLZF) and ROR-γt respectively, was biased toward the MAIT17 subset (PLZF^+^ROR-γt^+^) across most tissues, including the colon lamina propria (LP) as illustrated in Figure 1E. Notably, a significant fraction of liver MAIT cells fell into the MAIT1 category (PLZF^lo^ROR-γt^-^) (Figure 1E). Mirroring the phenotype seen in human MAIT cells, the majority of MAIT cells in CC011 mice either lacked or expressed low levels of the CD8 co-receptor (Figure 1F). Collectively, these findings confirm that the CC011 strain is rich in MAIT cells across both lymphoid and peripheral tissues, aligning more closely with MAIT cell frequencies typically found in humans^16–19^. Moreover, the MAIT cells from the CC011 strain predominantly exhibit an effector profile characteristic of the MAIT17 subset.

### Morphological and histological features of spontaneous colitis disease in CC011 emulate human pathology

The CC011 strain, which has previously been reported to display sporadic rectal prolapse, was further investigated and found to exhibit histological signs of colitis, even in the absence of any known murine pathogens^37^. Classified as a “Tier 1” strain by the University of North Carolina Systems Genetics Core Facility, CC011 is a strain that reflects a mosaic of all eight founder strains and is characterized by at least 90% homozygosity (https://csbio.unc.edu/CCstatus/index.py?run=AvailableLines). “Tier 1” indicates that these strains are well-established and widely used within the research community due to their genetic stability and representation.

Acknowledging that environmental factors significantly influence the onset of IBD, we confirmed that the spontaneous colitis phenotype was reproducible in our facility setting (Figure 2). The colon’s gross morphology in both sexes of the CC011 mice displayed signs of inflammation, particularly in the proximal colon, with some animals also showing inflammation in the mid and distal colon (Figure 2A). By contrast, the colons from the B6 control group exhibited no abnormal features. Notable observations in the CC011 mice included the presence of loose stool, a marked thickening of the colon wall, and discoloring (Figure 2A/B). Although these animals exhibited signs of colonic injury, they did not succumb to the disease. Additionally, we noted splenomegaly in older CC011 mice, which aligns with prior reports (Figure 2C)^37^. When comparing the ratio of spleen weight to body weight, there was a significant increase in the CC011 mice relative to the B6 controls, further supporting the manifestation of systemic inflammation in this strain (Figure 2C). Hematoxylin and eosin (H&E) staining of the CC011 colon sections demonstrated histopathological changes akin to those seen in human IBD. These changes included denuding of the surface epithelium and hypertrophy of the mucosa layer (Figure 1E). We observed lymphoid and neutrophil infiltrates spanning from the mucosal layer into the mucosal layer, through the muscularis mucosa, and into the submucosa (highlighted by white arrows in Figure 2E-F), and instances of gland herniation into the submucosa (indicated by a blue arrow in Figure 2F)^39^. Neutrophils (polymorphonuclear leukocytes or PMNs), a hallmark of acute inflammation, were noted both within the mucosal and submucosal layers and forming inside of crypt abscesses (Figure 2G, H). The inflammatory process in the CC011 mice was not limited to the mucosal layer; rather, it was transmural, permeating deeply into the muscularis layer, mirroring a Crohn’s disease-like pattern (Figure 2I). Chronic inflammation in IBD patients is a well-documented risk factor for gastrointestinal malignancies^40^. In the CC011 mice, areas of epithelial regeneration exhibited signs of dysplasia (Figure 2J), characterized by nuclear pleomorphism, excessive nuclear height, and crowded epithelial nuclei^40^. Disease onset in CC011 mice occurred between 22-23 weeks of age, with severity peaking and remaining stable from 27 to 48 weeks as indicated by the proximal inflammation scores (Figure 2K-L). In contrast, the B6 control group maintained a normal histological appearance in their colons, with no significant increase in inflammation as the mice aged (Figure 2K and L). In summary, our work successfully reproduces the spontaneous colitis phenotype previously reported by Rogala et al.^37^, while also delineating the histopathological progression of the disease with age in the CC011 strain, which mirrors features of human colitis, including Crohn’s-like transmural inflammation, and ulcerative colitis-like crypt abscesses.

**Figure 2:**
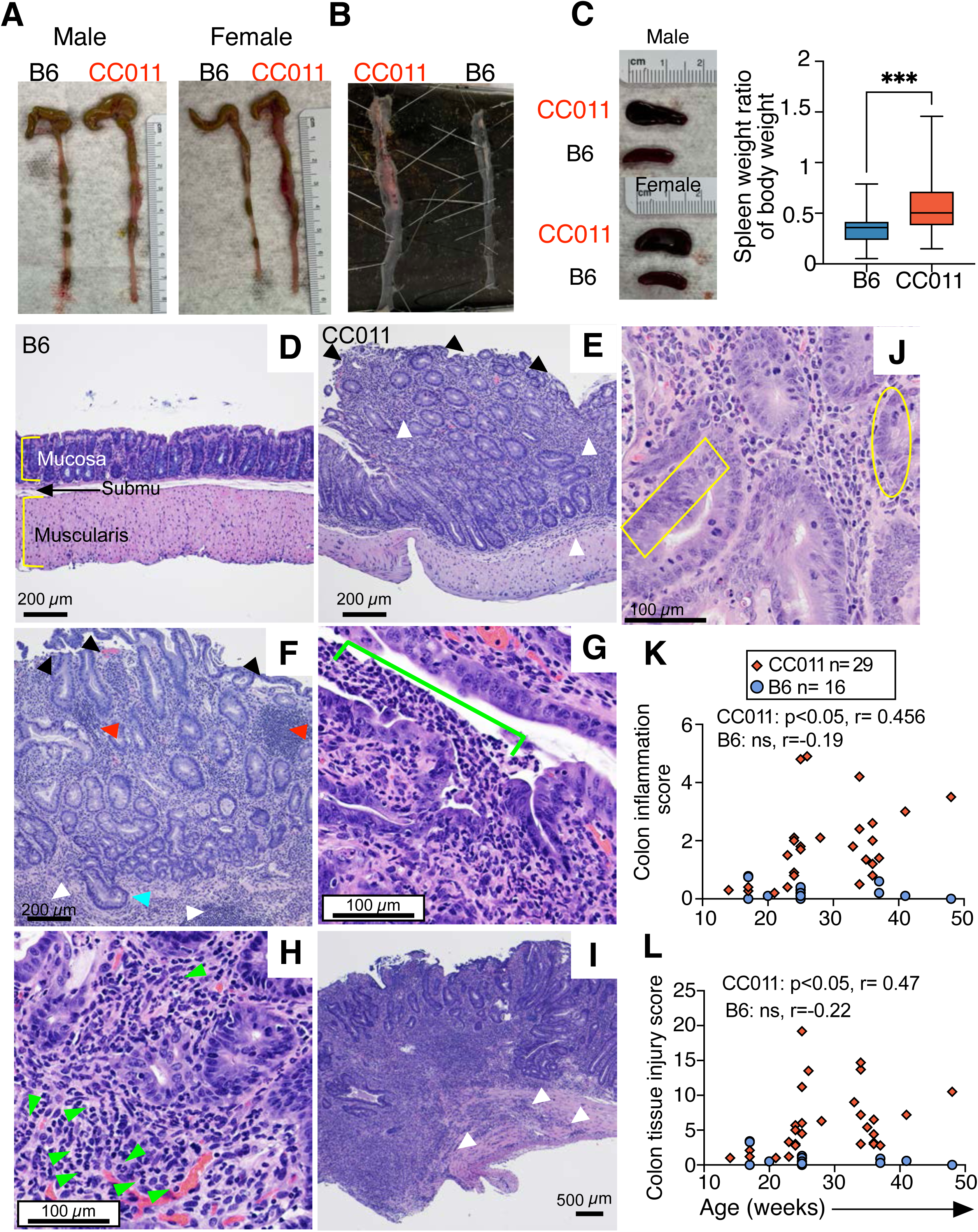
CC011 develop spontaneous colitis disease after 20 weeks of age with histological features that emulate human disease. A-B. Images of colons and C. spleens from B6 and CC011 mice week 27. C-H, Representative hematoxylin and eosin staining of paraffin embedded sections of colon from aged B6 (D) and CC011 (E-I) 27-36 weeks old. Black arrows denote denuding (striping) of the surface epithelium (E-F), white arrows denote inflammation in the mucosa and submucosa (E-F), red arrows denote lymphocytic accumulations, cyan arrow denotes gland herniating into the submucosa (F). G. Polymorphonuclear cells present in a crypt abscess of the mucosa, denoted by the fluorescent green bracket. H. Polymorphonuclear cells in the mucosa. I. Transmural inflammation in the muscularis, white arrows. J. Dysplasia of the colon epithelium (yellow oval and rectangle) in a week 35 CC011 mouse. K. Colon tissue inflammation and L. colon tissue injury scoring of H&E sections from CC011 and B6 mice on based on *Dieleman., et al*^59^ histological grading of colitis (\**P<.05*, Spearman rank correlation, (K, L) CC011 n=29, B6 n=16 mice/group).

### Colonic MAIT cells are the unconventional T cell subset that increases with age, and an imbalance of regulatory T cells is found in diseased CC011 mice

Given the age-dependent onset and increased severity of colitis, along with the substantial leukocytic infiltrates identified in H&E-stained sections of diseased colonic tissue (Figure 2), we aimed to unravel the dynamics of MAIT cells alongside other inflammatory and immunoregulatory cells in the colons of CC011 mice. To achieve this, we employed multiparameter spectral flow cytometry (Figure 3, with additional data in Supplementary Figure 1). Among the unconventional T cell subsets, MAIT cells (but not iNKT or γ8 T cells) showed an age-related increase within the lamina propria (LP) of the CC011 mice (Figure 3B, and Supplementary Figure 3). MAIT cells were also increased within the colon epithelium, interspersed between the epithelial cells (Supplementary Figure 4A). As established earlier, CC011 is a MAIT cell-rich strain under normal conditions. Intriguingly, in various extraintestinal tissues, we detected shifts in MAIT cell distribution that correlated with the age of the animals. Notably, in the liver, which is directly connected to the gastrointestinal tract, there was a discernible augmentation of MAIT cells concomitant with aging and disease progression (Supplementary Figure 4B). In contrast, in the lungs, we observed a reduction in MAIT cell numbers with age/disease, which might suggest a migration of MAIT cells to the colon from other tissue reservoirs (Supplementary Figure 4C). No age-related alterations were observed in the lymphoid tissues (Supplementary Figure 4D/E). The dysregulation or absence of regulatory T (Treg) cells is known to foster the onset of spontaneous colitis in animal models^2^. Moreover, maintaining a dynamic equilibrium between Treg cells and Th17 cells is crucial for curtailing inflammatory diseases^2^. Within the colon LP of CC011 mice, we observed an age-dependent increase in both Treg and Th17 cell populations (Figure 3C/D). Notwithstanding this increase, there was a concomitant decline in the Treg to Th17 cell ratio (Figure 3E), suggesting that while Treg cells are increasing, the Th17 cell population might be expanding at a faster rate. This imbalance may potentially impair the immunosuppressive function of Treg cells. To assess the suppressive capacity of CC011-derived Treg cells on the proliferation of conventional T (T_CONV_) cells following TCR activation, we performed co-culture experiments. CC011 Treg cells were mixed with Cell Trace Violet-labeled T_CONV_ cells from the spleen at varying Treg:T_CONV_ ratios, and the cells were stimulated with anti-CD3/CD28 beads (Supplementary Figure 5). The Treg cells from CC011 mice were proficient in suppressing the proliferation of activated T_CONV_ cells, demonstrating a comparable suppressive capability to their B6 counterparts, hence confirming that Treg cells from CC011 are functionally competent in modulating T cell activation during disease progression. However, the observed simultaneous rise of both Th17 and MAIT cells within the colonic environment suggests that in the context of the disease in CC011 mice, the Treg cell-mediated suppression may be insufficient. This raises the possibility that *in vivo*, the regulatory capabilities of Treg cells might be overpowered by the collective pro-inflammatory milieu created by the expanding Th17 and MAIT cell populations.

**Figure 3:**
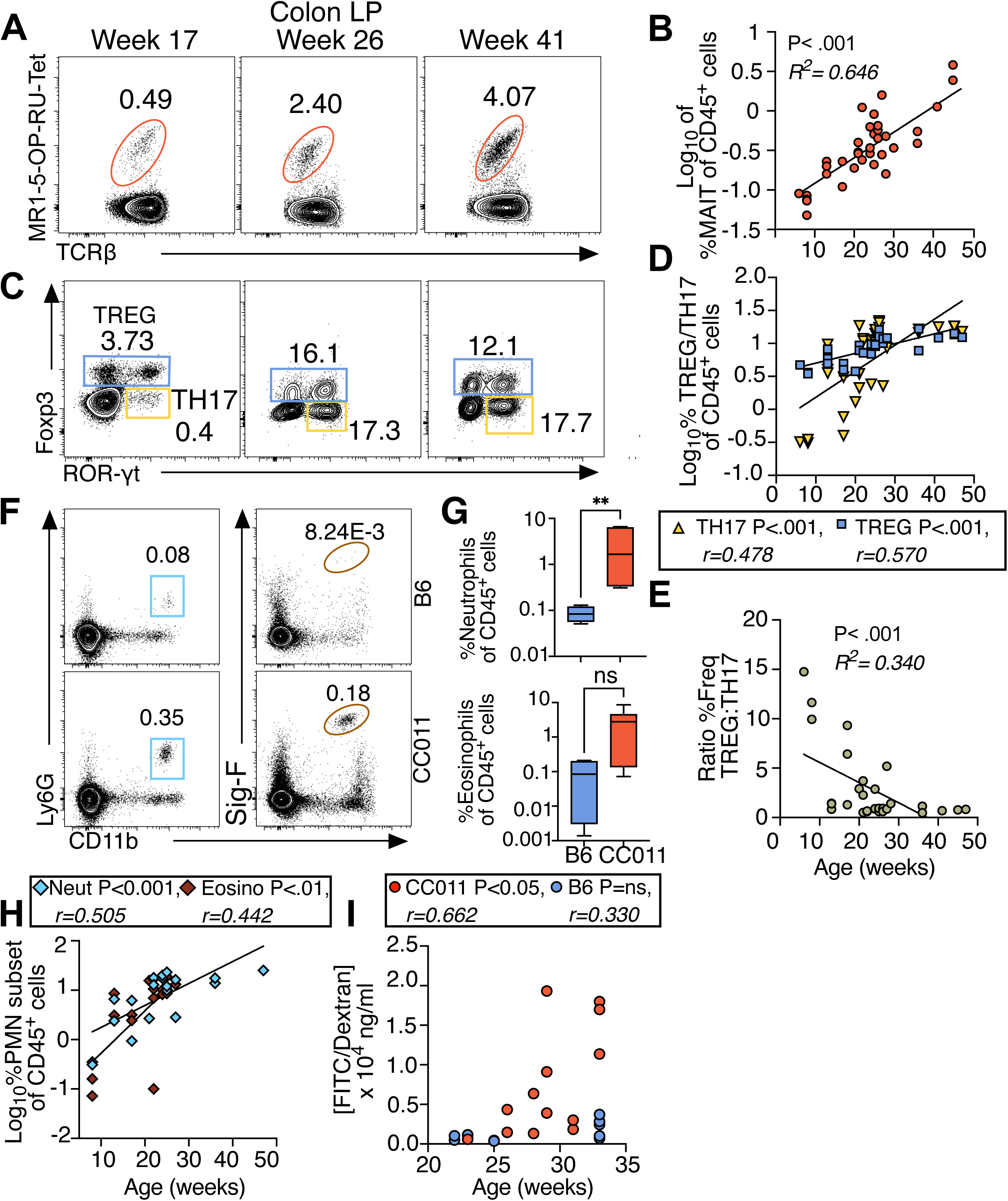
MAIT cells are the unconventional T cell subset associated with colitis disease onset. Representative contour flow cytometry plots of A. MAIT cells, C. Treg (CD4^+^Foxp3^+^ROR-γt^+/-^), D. TH17 (CD4^+^Foxp3^-^ROR-γt^+^) cells, and E. Neutrophils (CD11b+Ly6G^+^) isolated from the colon lamina propria (LP) of young and aged CC011 mice. Shown is the frequency of each cell subset of total live CD45^+^ cells. Log_10_ transformed data of B. MAIT (Red circles), D. TREG (Blue squares) /TH17 (Yellow triangles) of CD45^+^ cells and plotted against age (weeks). E. Ratio of the %Freq of Treg:Th17 cells. F. Representative contour flow cytometry plots of % frequencies of neutrophils and eosinophils (of CD45^+^ cells) in B6 and CC011 mice (8 weeks old) and summarized in G. in mice aged 8-17 weeks old, prior to disease onset. H. Log_10_ transformed data of neutrophil and eosinophil % frequencies (of CD45^+^ cells) from CC011 mice aged 8-47 weeks. H. FITC dextran concentration (ng/ml) in serum of orally gavaged CC011 or B6 mice. (*\*\*P<.01, ***P<.001*, Linear regression, (B) n=39; (D) Treg and Th17 n=33; (E) n= 33; (H) neutrophils n=20, eosinophils n= 16; Mann-Whitney Test, (G) neutrophils/eosinophils B6, n=4 CC011 n= 6 mice/group; Spearman rank correlation, (I) n=13; CC011, B6 n=12 mice/group).

Our histopathological analyses revealed the presence of neutrophils in CC011 colonic tissues using H&E staining at high magnification (Figure 2G, H). Further characterization through spectral flow cytometry revealed both increased neutrophils and eosinophils (Figure 3F). Notably, even in the absence of overt disease, CC011 mice displayed an approximately 200-fold increase in neutrophils within the colon compared to age-matched B6 controls, with a similar although less marked trend observed for eosinophils (Figure 3G). The temporal infiltration of these cells aligns with the progression of disease, with both neutrophils and eosinophils accumulating in the colon as the animals aged (Figure 3H).

The interplay between the immune system and commensal microbiota is intricate, with neutrophils, MAIT cells, Tregs, and Th17 cells all reacting to microbial constituents. Inflammatory processes within the gastrointestinal tract can disrupt the integrity of the epithelial barrier, a phenomenon commonly associated with IBD^41^. To assess this, we measured intestinal permeability in CC011 mice using fluorescein isothiocyanate (FITC)-Dextran passage from the gut into the serum following oral administration. At week 23, we found no significant difference in permeability between CC011 and B6 mice; however, an age-related increase in gut permeability was evident in CC011 mice as the disease progressed (Figure 3I). These data collectively indicate an active recruitment of innate immune cells to the disease site, which may be perpetuated by a compromised intestinal barrier, thus exacerbating the disease’s severity. Furthermore, a disproportionate ratio of Treg cells to the combined Th17/MAIT cell populations may also be a driving factor in the immunopathogenesis observed in this spontaneous model of colitis.

### Colonic MAIT cells from diseased CC011 animals produce IL-17A in vivo and display transcriptomic signatures consistent with Th17-differentiation, wound repair, and stimulation in a chronic inflammatory environment

Recognizing the elevated number of MAIT cells in the colon of CC011 mice in conjunction with the development of spontaneous colitis, and their potentiated MAIT17 effector phenotype, we aimed to delve into the transcriptional profile of these cells within the colon. The rarity of MAIT cells in common mouse strains poses a challenge for their isolation, particularly in B6 mice, hence we opted to compare the transcriptomic data of MAIT cells from the colon LP of diseased CC011 mice with those from the spleen of the same animals. We conducted single-cell RNA sequencing using the BD Rhapsody system, a droplet-based platform. We isolated MAIT cells from both the colon LP and the spleen by flow-sorting live MR1-5-OP-RU-Tet+TCRβ+ cells, followed by the generation of cDNA libraries for sequencing (Figure 4A). A total of 3,217 cells passed quality control and were further used for analysis. To identify and characterize subpopulation structures, we used unsupervised graph-based clustering, which led to the assignment of 8 distinct clusters (Figure 4B). Distinct niches were identified, primarily separating into colon (clusters 0, 2, 3, 6) and spleen (clusters 1, 4, 5, 7), indicating distinct transcriptional states (Figure 4C). Moreover, each individual animal contributed an even distribution of cells to each cluster, which is indicative of the representative tissue type (Figure 4D).

**Figure 4:**
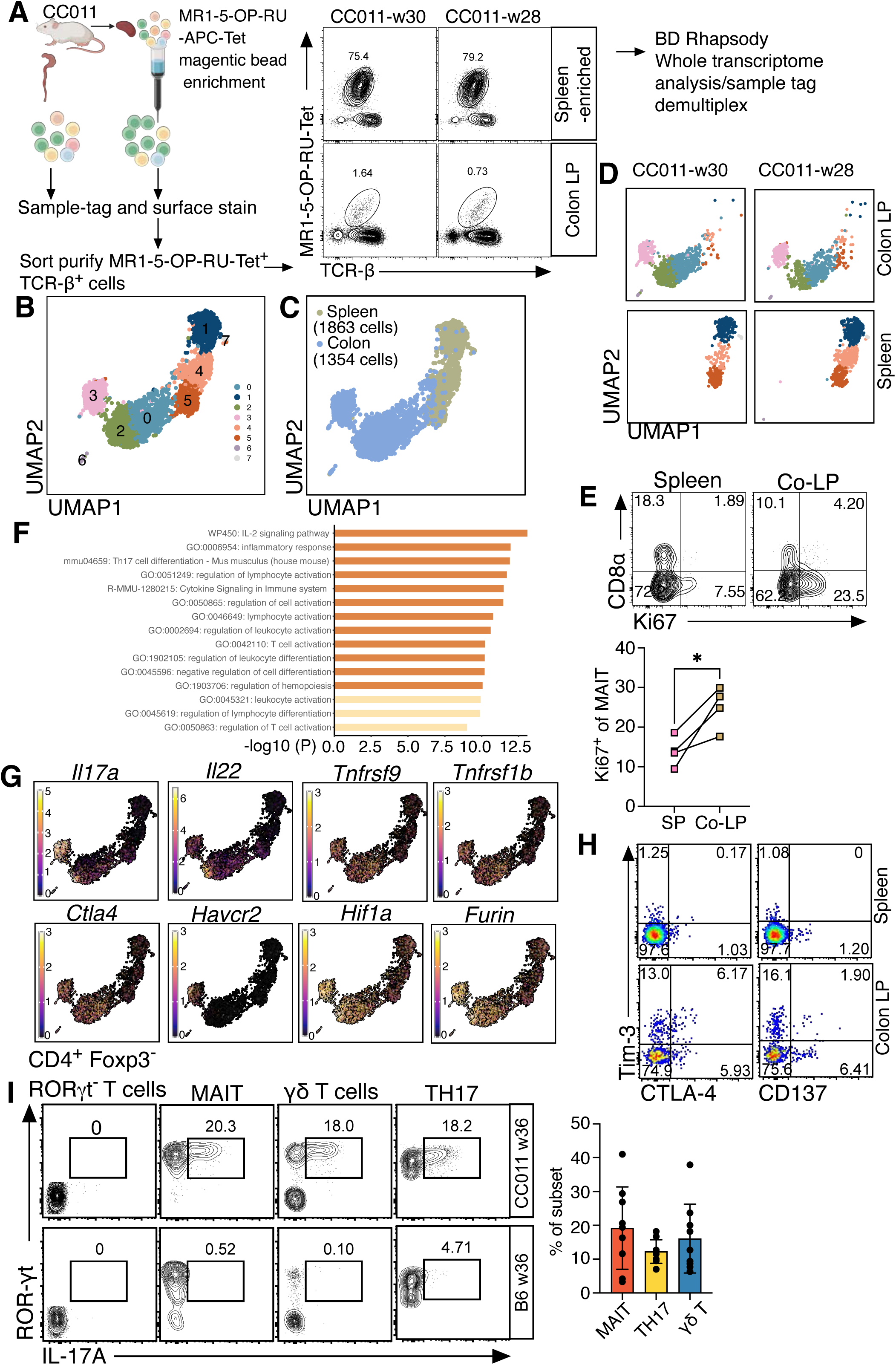
Colonic MAIT cells from diseased CC011 animals display a type III transcriptomic signature and hallmarks of activation in a chronic inflammatory environment. A. Experimental workflow for scRNAseq of MAIT cells isolated from spleen or colon LP of aged CC011 mice aged 28 and 30 weeks. B. Uniform Manifold Approximation and Projection (UMAP) representation of spleen and colon scRNAseq data, colored by Seurat clusters, C. tissue identity or D. original identity. E. Representative contour FACS plots of CD8α and Ki67 (*Mki67*) staining of MAIT cells in the spleen and colon LP of CC011 mice and summarized adjacent. F. Enriched terms from a colon tissue gene signature derived by the “FindMarkers” function in Seurat. Differentially expressed (DE) genes between colon and spleen (Table 2) were loaded into Metascape ^64^ for gene annotation and functional enrichment. Shown are the top 15 enriched terms where the color denotes the p-value. G. Feature plots of indicated genes. H. Representative FACS pseudo-color dot plots of Tim-3 (*Havcr2*), CTLA-4 and CD137 (*Tnfrsf1b)* surface expression, %expression in MAIT cells from spleen and colon CC011 are shown. I. Intracellular expression of IL-17A in MAIT cells was determined by flow cytometry without further *ex vivo* stimulation. Representative contour plots from B6 and CC011 (left) and summary graph of IL-17A production in CC011 colon T cell subsets (right) is shown. (\**P<.05,* Paired t-Test, (E) n= 4 mice/group).

Using the top differentially expressed genes that define each cluster, along with insights from the transcriptional diversity that exists amongst MAIT subsets in the thymus^15, 42^, we identified that clusters 3 and 2 from the colon were characterized by the expression of genes typically associated with the MAIT17 subset, including Il17a and Il22. However, there were differences in the levels of gene expression between these clusters (Supplementary Figure 6). Intriguingly, genes linked to cytotoxic functions, such as Serpinb9 and Serpinb6b, were prominent in cluster 3. Cluster 6 stood out as a small group of proliferating cells within the colon, corroborated by higher intracellular Ki67 expression observed via flow cytometry (Figure 4E). In the spleen, cluster 1 exhibited strong expression of genes typically related to the MAIT1 phenotype (Xcl2, Ly6c2, Klrd1). Clusters 4 and 5 appeared to represent cells that had undergone recent TCR activation, as evidenced by the upregulation of transcription factors like Klf2, Egr1, and Jun. Furthermore, cluster 7 demonstrated a heightened expression of genes involved in type I interferon signaling (Ifit1, Ifit3, Mx1, and Slfn5). To delineate the MAIT17 and MAIT1 profiles more distinctly, we employed gene signature scoring based on previously published data from Legoux et al.^42^, focusing on thymic MAIT cells (Supplementary Table 3). This analysis pointed to a significant enrichment of the MAIT17 signature across all colon-derived clusters and clusters 4 and 5 from the spleen (Supplementary Figure 7A). Conversely, the MAIT1 signature was particularly enriched in cluster 1 within the spleen (Supplementary Figure 7A). Building on recent findings that tissue-resident MAIT cells at barrier sites, such as the skin, also exhibit a transcriptome associated with tissue repair^30, 31^, we scored our CC011 MAIT cells against a gene set derived from skin-resident non-classical MHC-I-restricted CD8+ T cells involved in tissue injury^43^ (Supplementary Table 4). This comparison revealed that colon-derived MAIT cells showed significant enrichment for a tissue/wound repair signature when contrasted with their splenic counterparts (Supplementary Figure 7B). These results underscore the multifaceted roles of MAIT cells within different tissue contexts, particularly their involvement in both inflammatory responses and tissue repair processes in the colon.

A colon-specific gene signature for MAIT cells within the CC011 strain was identified, which is detailed in (Supplementary Table 4). Enrichment analysis of these genes highlighted key pathways, including IL-2 signaling, Th17 cell differentiation, and a spectrum of processes associated with lymphocyte activation and cytokine production (Figure 4F). We profiled the expression of pivotal genes implicated in Th17 cell differentiation (e.g., Il17a, Il22, Hif1a) alongside genes involved in immune activation during chronic inflammation (such as Tnfsf9, Tnfrsf1b, Ctla4, Havcr2, Furin), with their expression patterns presented in (Figure 4G). The distinct expression of Il17a in cluster 0 and Il22 in cluster 2 of colon-derived MAIT cells implies the presence of potentially functionally distinct subsets within the MAIT cell population residing in the colon. In spleen-derived clusters 4 and 5, we observed an enrichment for the MAIT17 signature (Supplementary Figure 7A), yet there was a relatively low expression of Il17a transcripts (Figure 4G). This observation may imply that the tissue environment influences the terminal functionality of MAIT cells.

We confirmed the expression of several activation markers (Tim-3, CTLA4, CD137) on MAIT cells extracted from the LP of diseased colons, with minimal expression detected in matched spleen samples (Figure 4H). To determine whether MAIT cells in diseased CC011 were actively secreting IL-17A, we administered brefeldin A, an inhibitor of protein transport, intravenously to aged CC011 mice before euthanasia. Subsequent analysis revealed that MAIT cells from the CC011 colon LP were indeed potent producers of IL-17A, contrasting with their B6 counterparts, which showed no such activity (Figure 4I). Although there was variability in IL-17A production among CC011 MAIT cells, there appeared to be a trend towards a higher proportion of these cells secreting IL-17A when compared with Th17 and γ8 T cells (Figure 4I).

These results reveal that the transcriptional landscape of MAIT cells from diseased CC011 colons is oriented towards a Th17 phenotype, indicating an adaptation to or an amplification of their environment’s inflammatory nature. These cells are not only transcriptionally active and highly activated, reflecting the chronic inflammatory milieu in which they reside, but they also exhibit a capacity for IL-17A production within the affected tissue. Intriguingly, when compared to their splenic counterparts in diseased CC011 mice, MAIT cells localized in the colon also show an increased expression of genes related to tissue repair. Thus, these cells may also play a reparative role within the tissue context, suggesting that MAIT cells in the context of IBD could be exerting a bifunctional influence on disease progression—both as contributors to the pathogenic inflammatory processes and as participants in tissue healing and repair. This dual functionality adds a layer of complexity to understanding the roles of MAIT cells in colitis and may inform the development of therapeutic strategies that could modulate these cells’ activity for better disease management and treatment.

### MAIT cell deficient Traj33^−/−^ CC011 mice exhibit minimal colitis disease severity

To explore the functional relevance of MAIT cells in the spontaneous colitis observed in CC011 mice, we utilized the CRISPR/Cas9 system to knock out the TRAJ33 gene segment from the CC011 genome, leading to a strain largely deficient in MAIT cells, referred to as CC011-Traj33−/−. Spectral flow cytometry analyses confirmed a significant reduction of MAIT cells in both lymphoid and peripheral tissues of aged CC011-Traj33−/− (Traj33−/−) mice, compared to their wild-type (WT) counterparts (Figure 5A/B). Upon examining other T cell subsets, including iNKT, γ8 T, Treg, and Th17 cells, we observed no substantial changes in most subsets. Notable exceptions included an approximate 2.5-fold increase in iNKT cells in the lungs and a roughly 5-fold decrease in Th17 cells in the liver of Traj33−/− mice (Supplementary Figure 9A/D). These alterations may suggest a potential interaction between MAIT cells and these populations in peripheral tissues. Most strikingly, aged Traj33−/− mice exhibited a pronounced reduction in colonic disease (Figure 5C-F). The colons of Traj33−/− mice lacked the thickening typical of diseased CC011 mice, and their stool was well-formed (Figure 5C/D). There was significantly less inflammation throughout the proximal and mid-colon (Figure 5D/E), with an absence of the lymphocytic aggregates that are usually present (Figure 5C). The mucosa showed minimal inflammation, with virtually no involvement of the submucosa (Figure 5C). Contrary to the CC011 WT mice (Figure 2), there was no evidence of dysplasia in the colons of the MAIT cell-deficient mice (Figure 5C). Corresponding with the reduced histological inflammation, myeloperoxidase immunohistochemistry, indicative of neutrophil infiltration, showed intense staining in the mucosa, submucosa and muscularis of WT CC011 mice, whereas MAIT cell-deficient mice displayed sparse positive staining, restricted primarily to the mucosa, revealing a significant reduction in neutrophil presence (Figure 5G, H). A similar reduction in neutrophil numbers was observed in the spleen and lungs and showed a trend in the liver of Traj33−/− mice (Figure 5I), which suggests that MAIT cells might influence neutrophil dynamics through immunological crosstalk. Overall, these findings implicate MAIT cells as critical players in the initiation and maintenance of inflammation in CC011 mice, serving a pathogenic role in the progression of colitis in this *in vivo* model.

**Figure 5:**
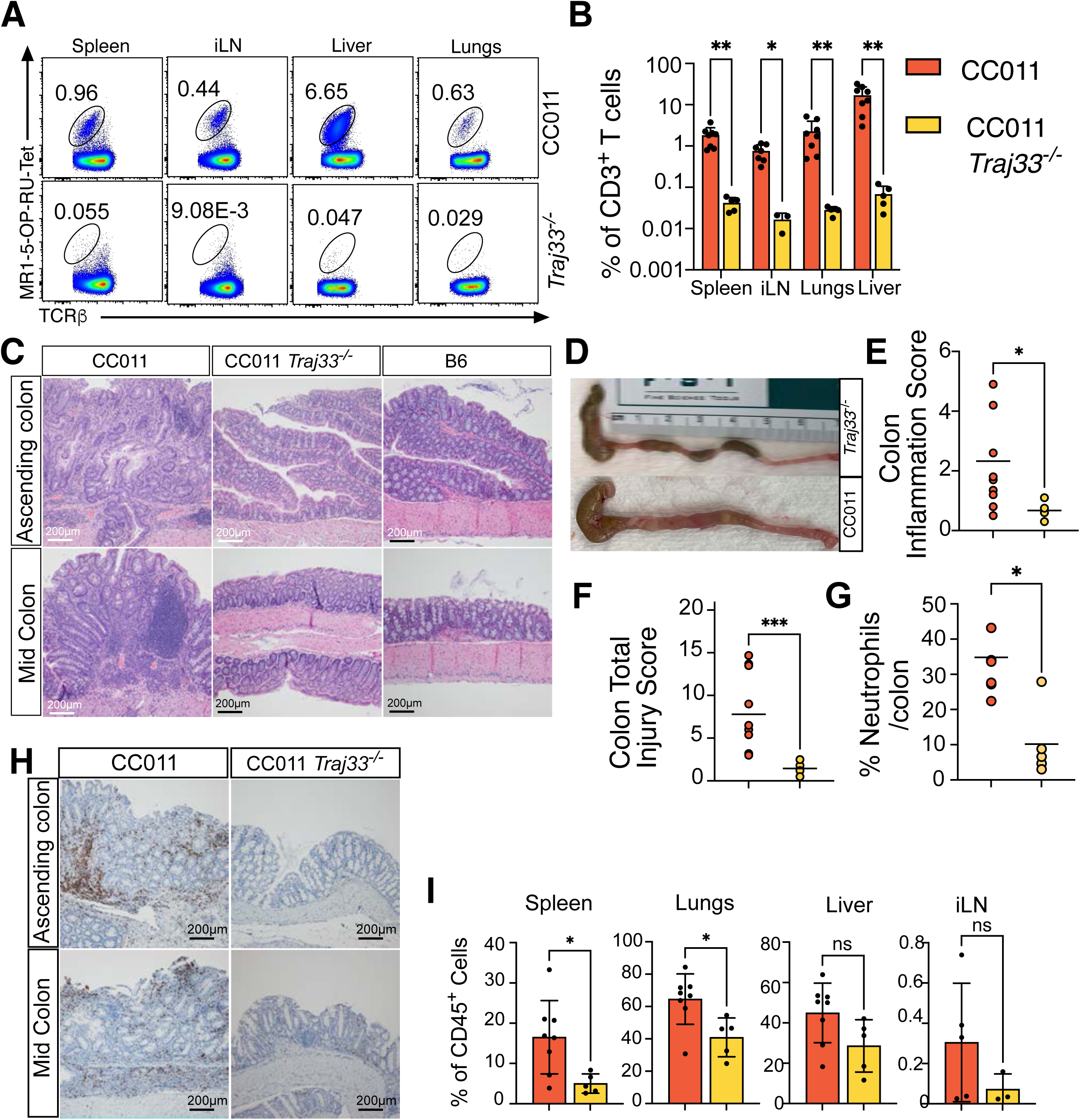
Limited colonic pathology and infiltration of neutrophils in the MAIT cell deficient CC011 Traj33^−/−^ strain. A. Representative pseudo-color plots of MAIT cells stained with MR1-5-OP-RU tetramer and TCR-β from different tissues isolated from CC011 and CC011-*Traj33^−/−^ (Traj33^−/−^)* mice 25-37 weeks old. The % MAIT cell frequencies of CD3^+^ T cells is shown and summarized in B. C. Representative hematoxylin and eosin staining of paraffin embedded sections of proximal (upper) and mid (lower) colon from aged CC011, *Traj33^−/−^* and B6 mice 25-37 weeks old. D. Images of colon showing, colon wall thickening and stool formation. E. Colon tissue inflammation and F. colon tissue injury scoring of H&E sections from CC011 and *Traj33^−/−^* mice on based on *Dieleman., et al* histological grading of colitis^59^. G. Representative myeloperoxidase immunodetection from paraffin embedded colon tissue sections derived from CC011 and *Traj33^−/−^* mice and H. summarized neutrophil counts/colon. I. %Neutrophil frequencies (of CD45^+^ cells) derived from FACS analyses of peripheral tissues from CC011 and *Traj33^−/−^*mice. (\**P<.05*, *\*\*P<.01*, Mann-Whitney (B) n= 3-7 mice/group, (D-G) n=5-7 mice/group, (H) n=3-7 mice/group).

## Discussion

The role of IL-17-producing lymphocytes in inflammatory diseases, including their immune-mediated effects, remains a contentious and context-specific topic. In this study, we have provided evidence within the CC011 strain that a high baseline presence of MAIT cells correlates with the progression of spontaneous colitis. Notably, the absence of MAIT cells in this strain resulted in a significant alleviation of the disease, suggesting a potential pathogenic role for these cells in this particular model of colitis.

Recent genome-wide association studies (GWAS) and meta-analyses have identified upwards of 240 susceptibility loci for Crohn’s disease and ulcerative colitis^32^. The numbers of loci grow when considering diverse ancestries^44^, pointing to a polygenic basis for IBD. In order to dissect the immunological underpinnings of IBD, common animal models have involved chemical induction in genetically predisposed strains^34^. The CC011 strain has been identified as having a genetic susceptibility to spontaneous colitis^37^, with quantitative trait loci (QTL) analysis attributing 27.7% of the phenotypic variance to genes located on chromosomes 12, 14, 1, and 8, suggesting that the disease in CC011 is likely governed by multiple genetic factors. This makes the CC011 strain a pertinent model for studying the complex genetics of human IBD and developing future treatments. The reproducibility of colitis in a different facility underscores a genetic etiology, although this does not exclude the possible influence of environmental factors on disease manifestation.

Compared to the B6 strain, which is resistant to colitis, the CC011 strain exhibits innate and innate-like immune subset alterations associated with inflammation, specifically implicating neutrophils and MAIT cells, even before the onset of colon pathology. For instance, the use of the CC011 strain in a model of severe asthma revealed that chronic exposure to house dust mite allergens led to significant airway remodeling and a high mortality rate^45^, characterized by Th2-mediated eosinophilia, which is distinct from the classical BALB/cJ strain response, reinforcing the susceptibility of CC011 to inflammation-driven diseases.

Across the 58 strains analyzed in this study, the higher baseline frequency of MAIT cells in the CC011 strain raises the question of whether this trait is under genetic control. Previous research, before MR1 tetramer technology, linked the abundance of MAIT cells to the TCRα locus on chromosome 14 ^46^ in the genetically related wild-derived inbred strain CAST/EiJ, which interestingly was not implicated in the QTLs associated with colitis phenotype in CC011^37^. Future investigations using expression QTL (eQTL) mapping could shed light on the genetic determinants of MAIT cell prevalence and their subsequent influence on disease in the CC011 strain. These insights could have far-reaching implications for understanding the genetics of IBD and for developing targeted interventions.

The contribution of MAIT cells to the pathology of colitis remains a topic of debate, with evidence pointing to both protective and pathogenic roles. Aligning with previous findings^28^, our study suggests a pathogenic involvement of MAIT cells in chronic colitis. The prominent Th17 transcriptional profile observed in the colonic MAIT cells, coupled with their robust production of IL-17A *in vivo*, positions them as significant contributors to inflammatory processes. From studies of ulcerative colitis patients, MAIT cells are reduced in the blood and also secrete more IL-17 compared to healthy controls^25^. Furthermore, at the site of colonic inflammation in UC patients the activated profile of MAIT cells reflects disease activity^25^. Our data further supports a pathogenic role for these cells via the observation that the absence of MAIT cells in CC011 mice correlates with a marked reduction in inflammation and overall disease severity. Taken together, our work supports that targeting MAIT cells may improve the severity of disease.

The potential interaction between granulocytes/neutrophils and MAIT cells in the context of colitis is particularly noteworthy. Neutrophil recruitment and activation can be prompted by granulopoiesis factors that are induced by IL-17A^1^. In the setting of IBD, chronic inflammation may extend the lifespan of neutrophils due to diminished apoptosis^47, 48^. Additionally, *in vitro* studies involving human cells have shown that MAIT cells, once activated by TCR engagement and cytokine stimulation, can modulate neutrophil survival through the production of TNF^49^. These findings underscore the complexity of the immunological landscape in IBD and highlight the necessity for further research to unravel the mechanisms of MAIT cell and neutrophil interaction. Understanding this interplay could lead to novel therapeutic targets and strategies, potentially allowing for more precise modulation of the immune response in IBD and related inflammatory conditions. The discovery that colonic MAIT cells in CC011 mice with colitis present a gene expression signature indicative of wound healing raises intriguing questions about their functional duality in IBD. The differential expression of Il17a and Il22, cytokines known for their roles in inflammation and tissue repair respectively, suggests that MAIT cells may embody a paradoxical role in IBD, contributing to both tissue damage and repair. This concept aligns with recent findings that skin-resident MAIT cells can promote wound healing^30, 31^, although similar in vivo studies in the gastrointestinal tract are yet to be conducted. Nevertheless, human studies have shown that activated MAIT cells from the blood can facilitate the closure of wounds *in vitro*, particularly when interacting with colonic epithelial cells, such as the CaCo2 cell line^50^. This capacity for wound closure highlights a reparative potential that MAIT cells could possess within the gut environment.

In the gastrointestinal tract, there’s an established dichotomy within the Th17 cell population, where both pathogenic and protective subsets coexist and are modulated by diverse microbial signals, leading to distinct metabolic and transcriptional profiles^51^. Similarly, MAIT cells may also exhibit such functional plasticity in the gut, being able to exert both protective and pathogenic effects depending on the context. In the CC011 model of spontaneous colitis, while the pathogenic aspect of MAIT cell function seems to predominate, their potential role in tissue repair should not be discounted. Further in-depth studies are necessary to dissect the conditions under which MAIT cells switch between these roles and how this balance can be therapeutically modulated to alleviate disease symptoms while promoting mucosal healing in IBD.

The intricate interplay between host genetics, the microbiota, and IBD is complex and multifaceted. Chronic inflammation associated with IBD often leads to an imbalance in the gut microbial community, known as dysbiosis^33, 52^, which can further exacerbate the disease state. Additionally, MAIT cells are known to be influenced by the commensal microbiota; they require bacterial antigens for their development and expansion in the periphery^14, 30, 53, 54^. In gnotobiotic (microbiota-free) environments, MAIT cells are less abundant compared to specific pathogen-free (SPF) conditions, indicating the significance of microbial signals for their homeostasis. The dysbiosis characteristic of chronic inflammation could potentially impact MAIT cell proliferation and function. The increased permeability of the intestinal epithelium observed in diseased CC011 mice could allow translocation of bacteria or bacterial products, which might stimulate MAIT cells, contributing to their activation and expansion during disease. It remains an open question whether the microbiome of CC011 mice harbors unique microbial communities that not only support the basal development of MAIT cells but also the progression of colitis. Experimental models wherein microbiota from different sources are introduced to gnotobiotic mice have demonstrated that microbiome composition can significantly influence colitis outcomes, even providing protective effects depending on the microbial consortia introduced^55, 56^. Future investigations could elucidate the role of the microbiota in MAIT cell-driven colitis through interventions such as antibiotic treatments or fecal microbiota transplants in germ-free CC011 mice.

In summary, our study adds to the growing body of *in vivo* evidence indicating that MAIT cells can have a pathogenic role in spontaneous colitis. Targeting MAIT cells with novel inhibitors might offer a supplementary approach to treating active IBD, leveraging the CC011 strain and related resources. Additionally, monitoring MAIT cell activity could serve as a non-invasive biomarker for disease activity, potentially informing and guiding treatment decisions for IBD patients.

## Acknowledgements

The authors thank Ana Campos Codo, Neus Font Porterias and Rosemary Callahan for their technical assistance. E. Erin Smith (University of Colorado, Anschutz Medical Campus, Pathology Shared Resource, Research Histology Lab, in part supported by the Cancer Center Support Grant (P30CA046934), and Katie Tuscan (Cardiovascular pulmonary histology core) for immunohistochemical stain assistance. Tinalyn Kupfer, for flow sorting and managing the Flow Cytometry Facility (University of Colorado, Immunology and Microbiology). Lee Schuyler, at the Genomics Core, University of Colorado, Anschutz Medical Campus. Rachel Lynch for the co-ordination of the CC-strains (University of North Carolina, at Chapel Hill). The Mouse Genetics Core at National Jewish Health, for the generation of CC011-Traj33−/− mice.

## Methods and Materials

### Animals

CC011/Unc strain (CC011) were bred in house using CC011/Unc mice from the University of North Carolina at Chapel Hill (UNC). C57BL/6 were purchased from Jackson Laboratories (JAX) and housed at the University of Colorado Anschutz Medical campus until ready for use at the specified age. All mice used for this study were between 6 to 47 weeks of age and both male and female mice were analyzed. For other CC– and Founder-CC strains (Table S1) thymi were purchased from UNC and JAX respectively. Before their relocation to UNC, CC lines were generated and bred at Tel Aviv University in Israel, Geniad in Australia and Oak Ridge National Laboratory in the USA. The CC011-*Traj33^−/−^* strain were generated at the Mouse Genetics Core Facility at National Jewish Health using CRISPER/Cas9-mediated^57, 58^ deletion of the *Traj33* gene in CC011 zygotes. Two guide RNA motifs (Guide #268for CTGGAATCCATAATGGGCAAGGG; Guide #228rev TCATATATCGAAGACTTACCTGG) were used to eliminate the *Traj33* gene segment from the genome of CC011 mice (Figure S8). Zygotes were then transferred into pseudopregnant recipients. The pups were genotyped by PCR using primers (TRAJ33_268For-5: TGTATGAGAGCACAGTTCCTC; TRAJ33_156-228-3: AGTGGGATGGGCTGGCTTATC) outside the region to be modified to identify putative positive founders. The founders were tested for modification of the *Traj33* gene using NGS (Figure S8). All mice were raised in a specific pathogen-free environment at the Office of Laboratory Animal Research at the University of Colorado Anschutz Medical campus or National Jewish Health, Colorado. All animal procedures were approved by the UCD (00065) Institutional Animal Care and Use Committees and were performed in accordance with the approved guidelines at both facilities. At the specified age, colons were embedded and processed for histology. Hematoxylin and eosin-stained tissue was scored by a histo-pathologist blinded to the treatments and groupings of animals and using described methods^59^.

### Tissue preparation and cell suspensions

Single cell suspensions were prepared from the colon after removing fecal matter by flushing PBS though the colon via a blunt-end 16 gauge syringe. The epithelial layer was dissociated by shaking whole colon in IEL solution (RPMI, 2% FCS, 5% DTT, 1mM EDTA, 7.5mM HEPES) at 200rpm, 37°C for 15 mins. After dissociation of the epithelial layer, the supernatant was filtered through a 100-μm cell strainer and pelleted by spinning at 500 x g for 10 mins at room temperature. The IELs were resuspended in complete media (cRPMI) (RPMI containing 10% fetal calf serum and enriched supplements) prior to Percoll gradient separation. The remaining colon tissue was minced using scissors in digestion media (RPMI, 100 μg/mL of DNASE I (Sigma Aldrich), 0.5 mg/mL Collagenase type IV (Worthington Biochemical Company), 7.5mM HEPES, β-mercaptoethanol, 5% FCS). Colon tissue was digested for 25 mins at 37°C with shaking at 200 rpm. Digested tissue was filtered through a 100 µm cell strainer followed by centrifugation and resuspension in 40% Percoll prior to overlay on a 70% Percoll gradient. After centrifugation the middle cellular layer was harvested for flow cytometry. To prepare single-cell suspensions from lymphoid tissues, thymus, spleen or lymph nodes were mechanically dissociated by pressing each individual tissue with the back of a syringe plunger through a 70-μm nylon mesh strainer into cold cRPMI media. Splenocytes were subjected to red blood cell lysis before resuspension into FACS buffer (PBS, 0.5% BSA, 0.5mM EDTA, 1% Azide). Liver and lung single-cell suspensions lungs were prepared as previously described^60^.

### Flow Cytometry

Single cell suspensions were stained with efluor780 viability dye (ThermoFisher) for 10 mins at room temperature and washed once prior to cell surface staining. Cell suspensions were stained with a the MR1-5-OP-RU-APC tetramer, CD1d-PBS57-BV421 tetramer (kindly provided by the NIH Tetramer Core Facility), and a combination of cell surface markers (Table S2) in FACS buffer at room temperature for 20 mins. Cells were washed twice and fixed with Foxp3 Transcription Factor Staining Kit (ThermoFisher), according to the manufacture’s recommendations. Intracellular staining for transcription factors was performed on ice for 1.5hr. Subsequently cells were washed twice in perm/wash buffer and resuspended in FACS buffer prior to acquisition. Intracellular cytokine staining for IL-17A was performed in the same manner as transcription factors, animals were injected with BFA 3 hours before tissue harvest (200μl of PBS 0.25mg/ml delivered i.v/mouse). Data were acquired on the Cytek Aurora flow cytometry system using SpectroFlo software (v3.0) and further analyzed using FlowJo software v10.7.1 (BD Biosciences).

### Fluorescein isothiocyanate (FITC)-Dextran Flux

Mice were orally gavaged (0.6 mg/kg body weight) with 4 kDa dextran labeled with fluorescein (Sigma) and serum was collected in EDTA containing blood collection tubes (BD) prior to and 4 hours after oral gavage. The amount of fluorescence was measured with a fluorimeter (Tecan) at 490_exc_/525_em_ nm. A standard curve was generated to calculate the amount of dextran that was present in the serum. Pre-treatment serum fluorescence was subtracted from post treatment measurements to account for any background fluorescence of serum.

### T_reg_ mediated suppression of CD4^+^ T cell proliferation

An *in vitro* Treg suppression assay was developed based on a previous study ^61^. Splenic CD4^+^ T_conv_ cell and T_reg_ were isolated using the mouse CD4^+^CD25^+^ Regulatory T Cell Isolation Kit (Miltenyi) according to the manufacturer’s instructions. T_conv_ cells were labeled with 1μM of CellTrace Violet (ThermoFisher) according to the manufacturer’s instructions. In a 96 well U-Bottom plate, 1 x10^5^ T_conv_ cells were stimulated with mouse T-cell Activator CD3/CD28 Dynabeads (ThermoFisher) at a cell to bead ratio of 3:1 and co-cultured in a 37°C incubator (5% CO_2_) without or with T_reg_ cells at the specified ratios (Figure S5). After 3 days cultures were harvested and stained with viability dye and cell surface markers (Table S2) and subsequently fixed with 2% PFA. Samples were acquired on the Cytek Aurora flow cytometer to determine proliferation kinetics.

### Tetramer Enrichment of MAIT cells with MR1-5-OP-RU-PE-Tetramer

To enrich for splenic MAIT cells, up to 5×10^7^ cells were incubated with MR1-5-OP-RU-PE-Tetramer in MACS buffer (0.5% BSA, 2mM ETDA, PBS), for 25 mins at room temperature. Cells were washed twice and incubated with anti-PE microbeads (Miltenyi), followed by separation using an autoMACS Pro Separator (Miltenyi) according to manufacturer’s instructions. PE-microbead-labelled cells in the enriched fraction were stained with the full panel of antibodies.

### Single cell RNA sequencing

Single cell whole transcriptomes were prepared using the BD Rhapsody Single-Cell Analysis System (BD Biosciences) according to the manufacturer’s specifications. Prior to cell sorting on the Aria 3 (BD Biosciences) and during cell surface antibody staining, up to 2 x10^6^ MR1-5-OP-RU-APC Tetramer magnetic bead enriched spleen (n= 2) or unenriched colon (n= 2) single cell suspensions were labeled with an oligonucleotide-tagged antibody sample tag (BD Biosciences). MAIT cells (MR1-5-OP-RU-Tet^+^TCRβ^+^CD3^+^) were sorted after doublet, and viability discrimination (Figure 4). Prior to cDNA library preparation, all MAIT cells from the different animals and tissues were pooled, 4 unique sample tags were combined in the library. Libraries were sequenced on an Illumina NovaSeq platform at the University of Colorado Genomics Core with the following read lengths: read 1 – 150 cycles; read 2 – 150 cycles; and i7 index – 8 cycles.

### ScRNAseq Analysis

The quality of sequencing reads was evaluated using FastQC and MultiQC. Sequencing reads (FASTQ) were mapped and sample Tag deconvoluted with The BD Rhapsody™ WTA Analysis Pipeline on the GRCm38 genome sequence. This pipeline produced a gene expression matrix for each sample, which records the number of UMIs for each gene associated with each cell barcode. Aggregated data were then imported into the R environment and analyzed with Seurat (4.1.3). Low-quality cells were filtered using the cutoffs nFeature_RNA >= 500 & nFeature_RNA < 3000. Cells with a high mitochondrial content were removed using the percent.mt < 15 function in Seurat. Genes expressed in less than 20 cells were ignored. This resulted in 3217 cells with 13,875 genes for downstream analyses. Dimensionality reduction was performed prior to integration for visualization purposes, by selecting 2000 highly variable genes for principal component analysis (PCA) and uniform manifold approximation and projection (UMAP). The data were used in the subsequent nonlinear dimensional reduction with the RunUMAP function (20 dimensions, resolution 0.9). This dataset was subsequently split up into 4 samples from 2 different tissues and animals based on sample cell hashing tags/barcodes. Some of the plots displaying cells on UMAPs were generated using the SCpubr package^62^.

### Identification of differentially expressed genes between clusters and tissues

We identified cluster-enriched genes by using the FindAllMarkers function in Seurat with test.use = wilcox. This function identified differentially expressed genes for each cluster by comparing the gene expression for cells belonging to a cluster versus cells belonging to all other clusters. Only those genes that passed an adjusted p value (Benjamini-Hochberg) cutoff of 0.05, log fold change > 0.4 and min.pct = 0.2 were included in the downstream analyses. The top 5 marker genes for each cluster were displayed using the DotPlot function in Seurat. A colon gene signature was derived by calculating differential gene expression between the colon and spleen tissues using the FindMarkers function in Seurat with test.use = MAST^63^ and a minimum percentage of 20% and logfc threshold of 0.4. Unbiased analysis of the resulting gene list (Supplementary Table 4) was performed on the online resource Metascape^64^, where enriched biological pathways were calculated. The top 15 pathways were re-plotted using ggplot. To score for MAIT1 ^42^, MAIT17 ^42^ and wound healing^43^ signatures, previously published genes signatures were calculated on the Seurat object using the Addmodulescore function in Seurat and displayed using SCpubr feature plots.

### Myeloperoxidase staining

Four-micron thick paraffin sections were prepared for immunodetection of Myeloperoxidase (Abcam, Cambridge, MA; ab208670; Lot: GR3390666-1; clone EPR20257; 1:1500 dilution). Antigens were revealed in BORG solution (pH 9.5, Biocare Medical) for 10 min at 110°C (NxGen Decloaker, Biocare) with a 10 min ambient cool down. Immunodetection was performed on the Discovery Ultra stainer (Ventana/Roche) with primary incubations for 32 min at 37°C. Myeloperoxidase was visualized with the OmniMAP anti-Rabbit HRP and ChromoMAP DAB kit (Venata/Roche). All sections were counterstained in Harris hematoxylin for 2 min, blued in 1% ammonium hydroxide, dehydrated in graded alcohols, cleared in xylene and cover glass mounted using synthetic resin. Negative controls to confirm the specificity of the immunostaining included omission of the primary antibody incubation step in the IHC protocol, substitution of the primary antibody diluent.

### Statistical analyses

The statistical tests that were used include: Mann-Whitney, unpaired t-test, paired t test, simple linear regression, Spearman rank correlation. All tests were performed using Prism version 10 (GraphPad Software). Results were considered statistically significant when p < .05 and “ns” if the comparison was non-significant. For each experiment, number of independent experiments, replicates and the statistical tests used are indicated in the figure legends.

## Figure Legends

**Figure S1:**
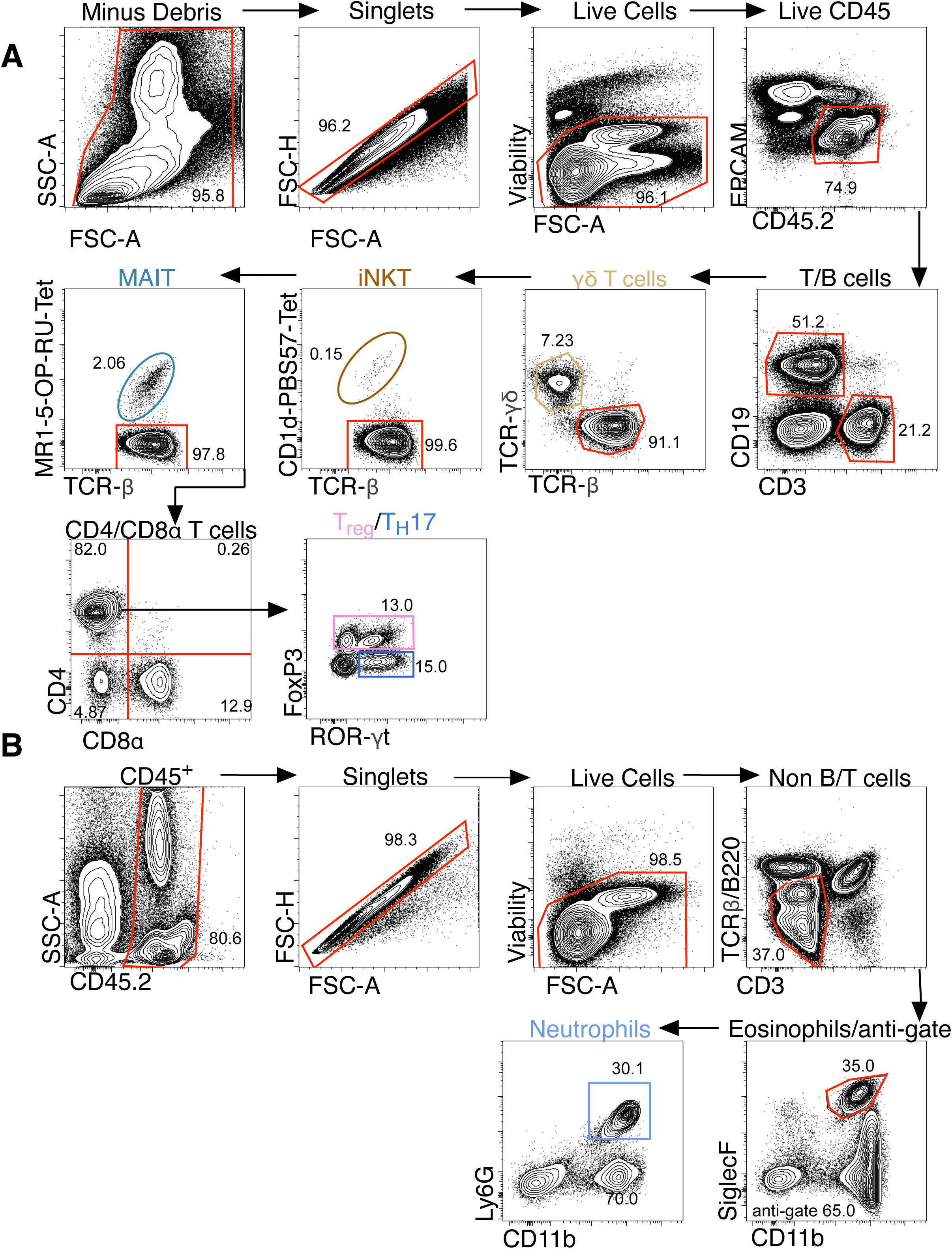
Cytek Aurora spectral flow cytometry gating strategies. A. Shown is the gating strategy for tissue T cells and B. Neutrophils. Frequencies of the parental gate are shown.

**Figure S2:**
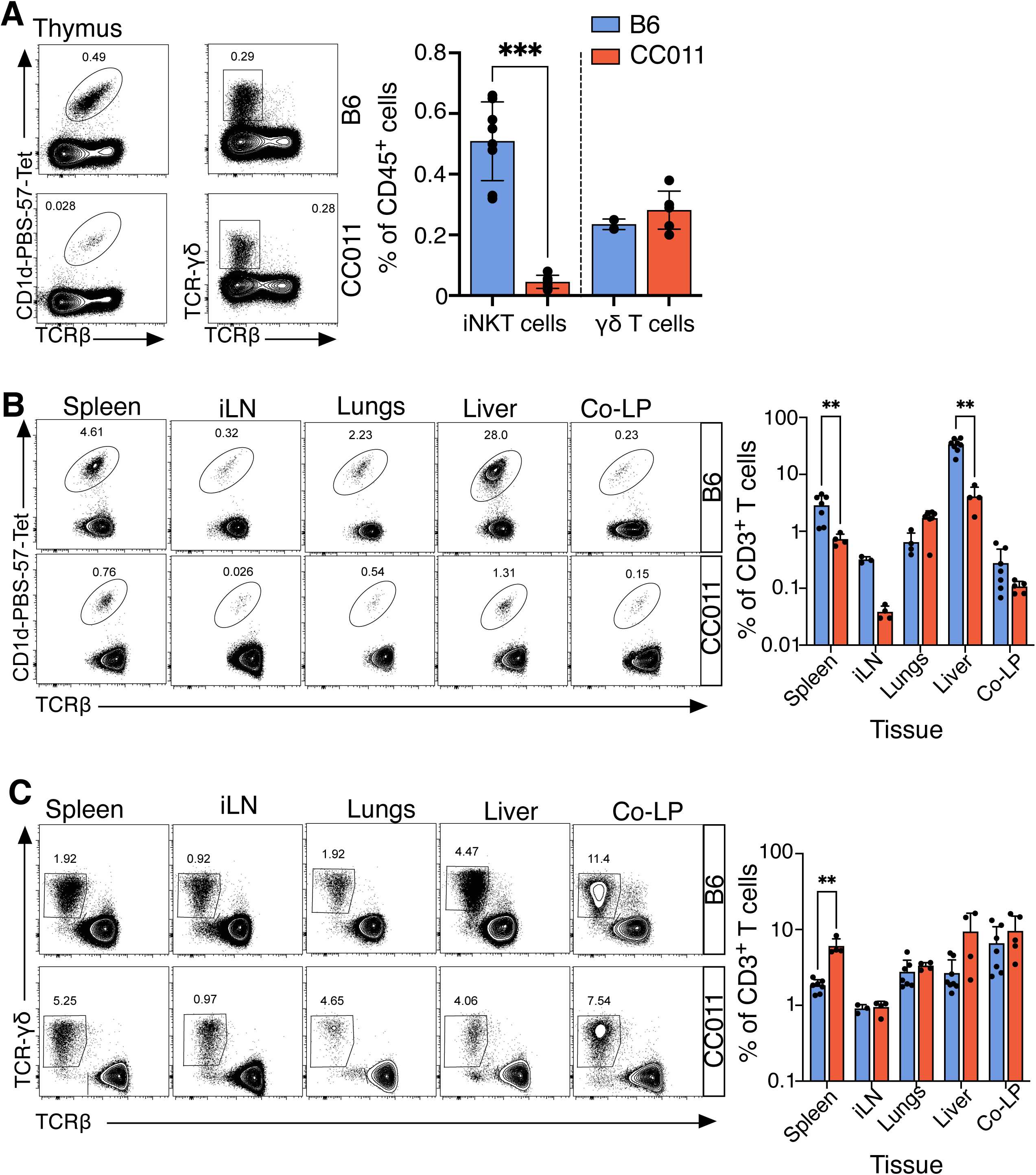
Young CC011 iNKT and γ8 T cell dynamics. A. Representative contour plots of iNKT and gd T cell staining in the thymus of CC011 (lower, blue) and B6 (upper, red) mice aged 6-9 weeks old. Show is the %frequency of each subset of CD45^+^ cells and summarized on the right panel. B. Show are representative contour plots of tissue iNKT and C. γ8 T cells aged 6-9 weeks old, and the %frequencies of each subset of CD3^+^ T cells are summarized on the right panel. (\**P<.05*, *\*\*P<.01*, *\*\*\*P<.001,* Mann-Whitney (A) n= 4-8 mice/group, (B, C) n= 3-8 mice/group).

**Figure S3:**
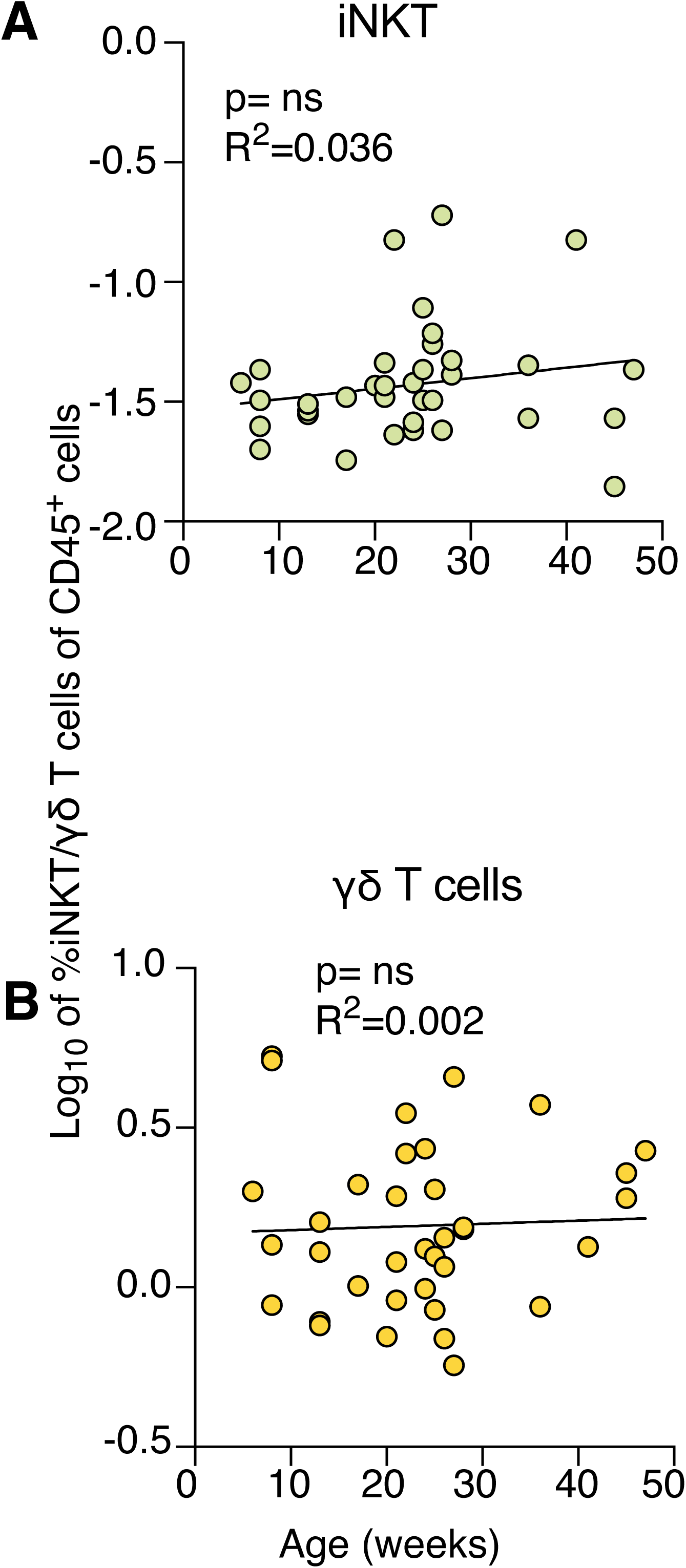
iNKT and γ8 T cell kinetics in the Co-LP. Shown are linear regression curves of Log_10_ transformed A. iNKT and B. γ8 T cell %frequencies (of CD45^+^ cells) as a function of age (weeks) in the colon-LP of the CC011 strain. (*P=ns*, Linear regression, (A) n= 37; (B) n= 37)

**Figure S4:**
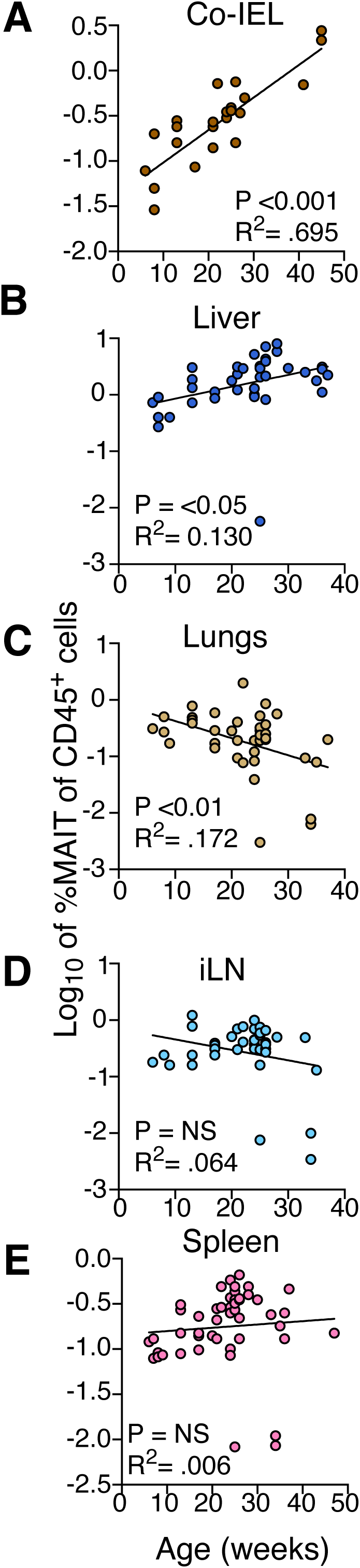
Tissue MAIT kinetics as a function of age. A-E. Shown are linear regression curves of Log_10_ transformed MAIT cell %frequencies (of CD45^+^ cells) as a function of age (weeks) across the different tissues in the CC011 strain. (\**P<.05*, **<0.01,*\*\*\*P<.001*, Linear regression, (A) n= 27, (B) n= 41, (C) n= 40, (D) n= 40 (E) n= 49)

**Figure S5:**
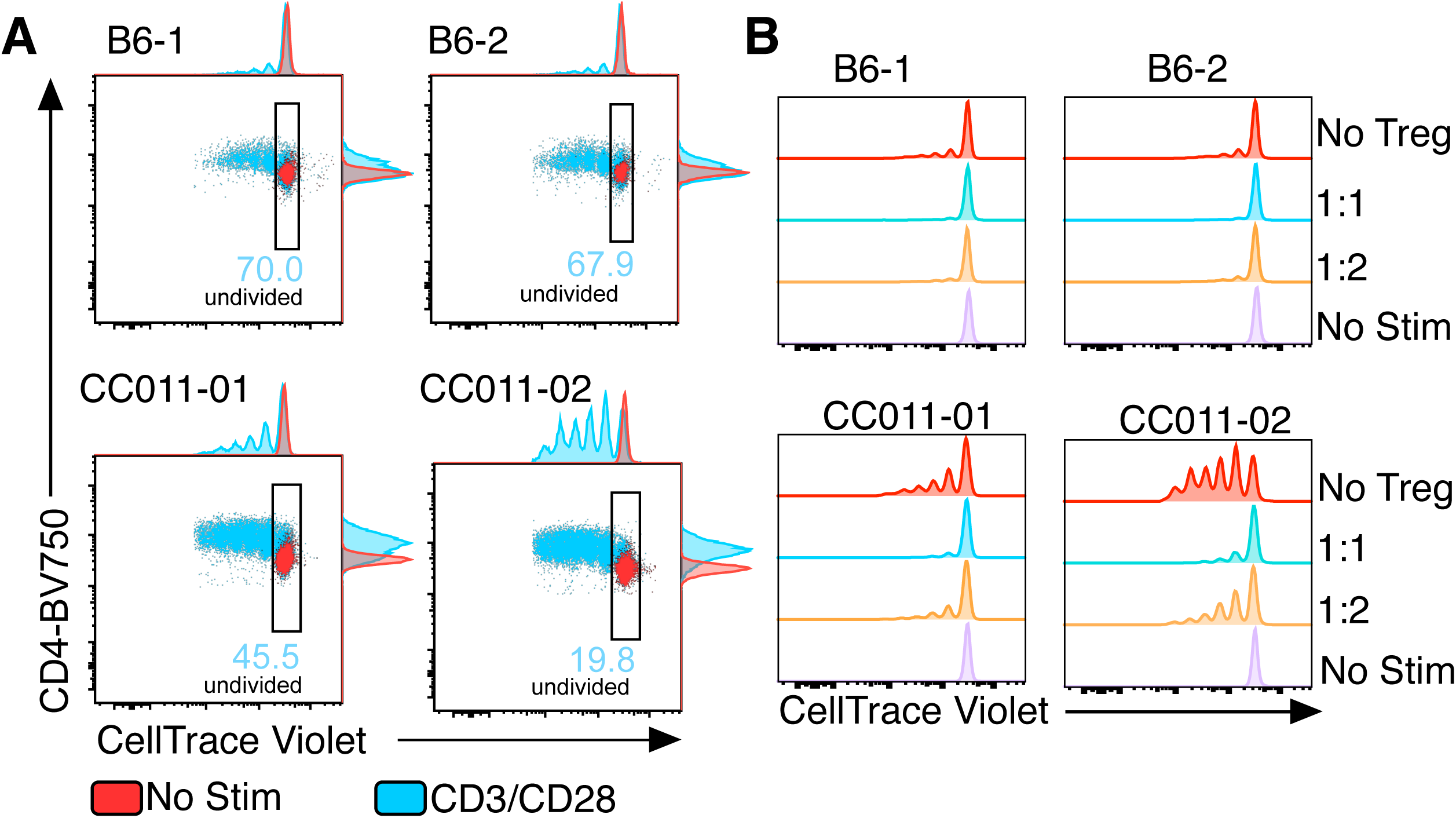
T_reg_-mediated suppression of T_conv_ proliferation. A. Shown are histograms/dot plots of CD4^+^ T_conv_ cell proliferation as measured by gating on CTV labeled T_conv_ from B6 (upper) and CC011 (lower) spleen aged 8 weeks old, after 3 days of culture at 37°C. Red depicts unstimulated T_conv_ cells and blue T_conv_ stimulated with a 3:1 (cell:bead) ratio of CD3/CD28 T cell activator dynabeads. The %undivided of CD3/CD28 stimulated cells is shown. B. Summary of offset histograms of CTV dilution (x-axis) gated on T_conv_, the ratios reflect T_reg_:T_conv_ in the presence of CD3/CD28 stimulation in B6 (upper) and CC011 (lower).

**Figure S6:**
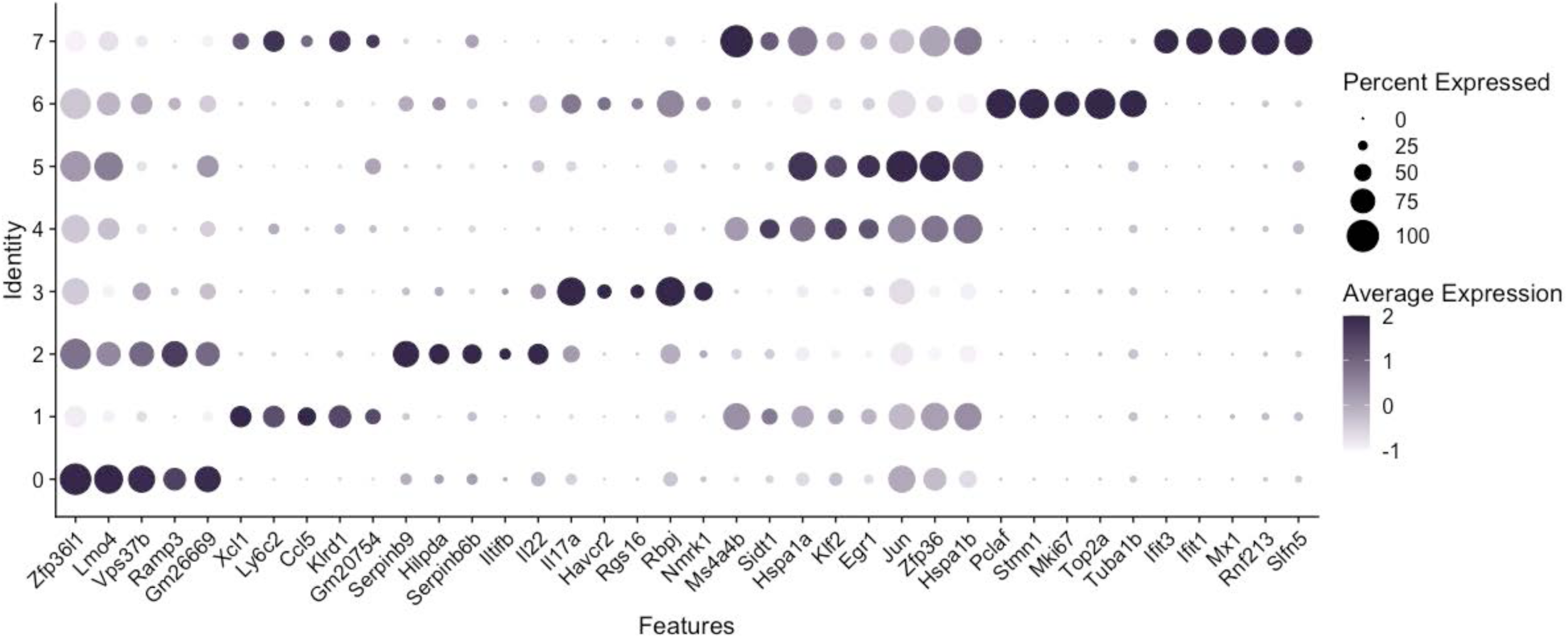
Top 5 genes/cluster scRNA seq. Shown is a dot plot of the Top 5 differentially expressed genes from each of the Seurat clusters 0-7. The gene identity was generated with the Seurat function FindAllMarkers. The size of the dot represents the percentage of cells of the cluster expressing the gene and the color gradient represents the relative expression value.

**Figure S7:**
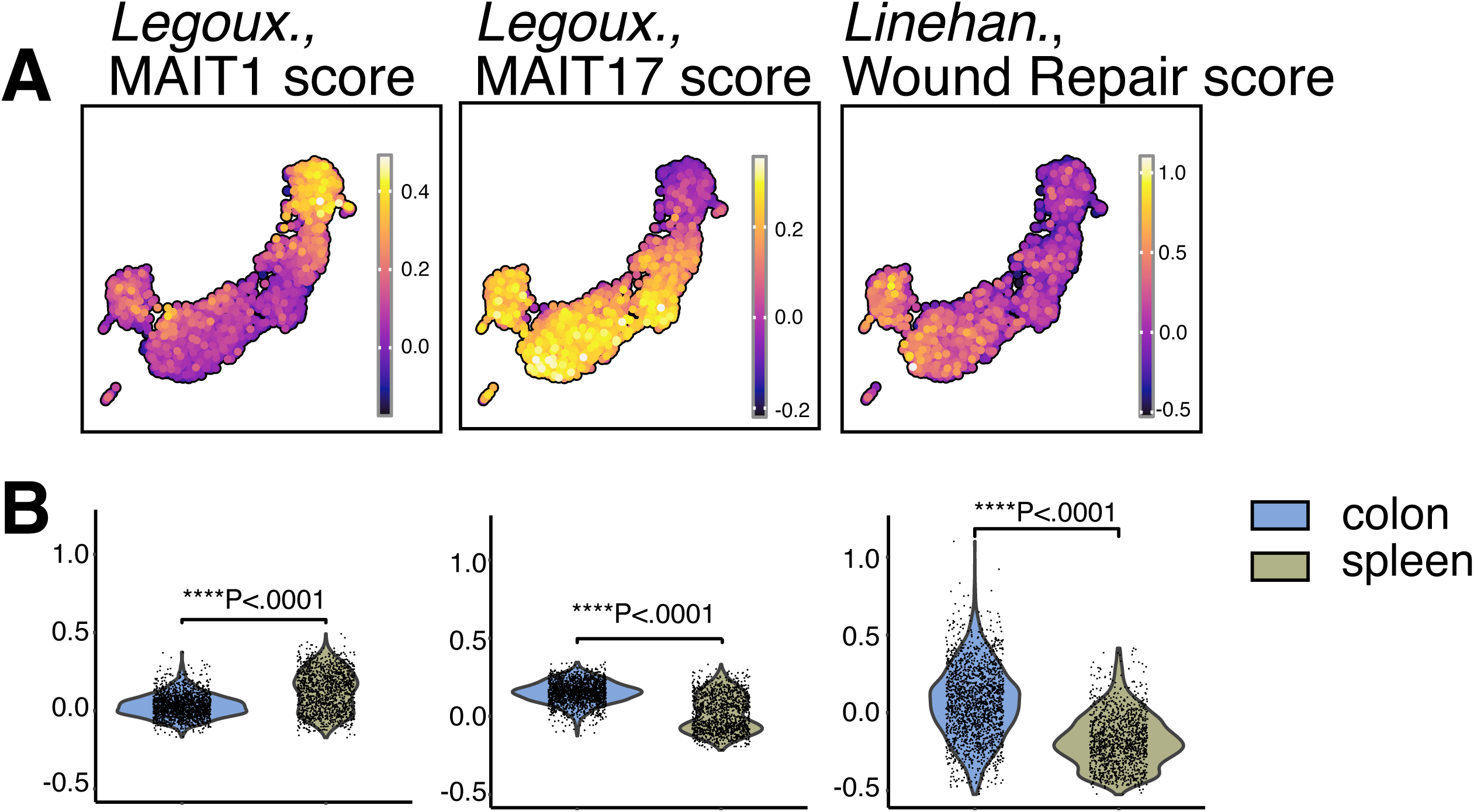
MAIT1, MAIT17 and wound healing scores. Shown are UMAP representations of A. MAIT1 and MAIT17 signature scores based on gene signatures from Legoux., 2019^42^ and wound repair scores are based on gene signatures from Linehan., et al^43^. B. Violin plots of signature score in A. separated by tissue. (***P<0.001, Wilcox Test (B))

**Figure S8:**
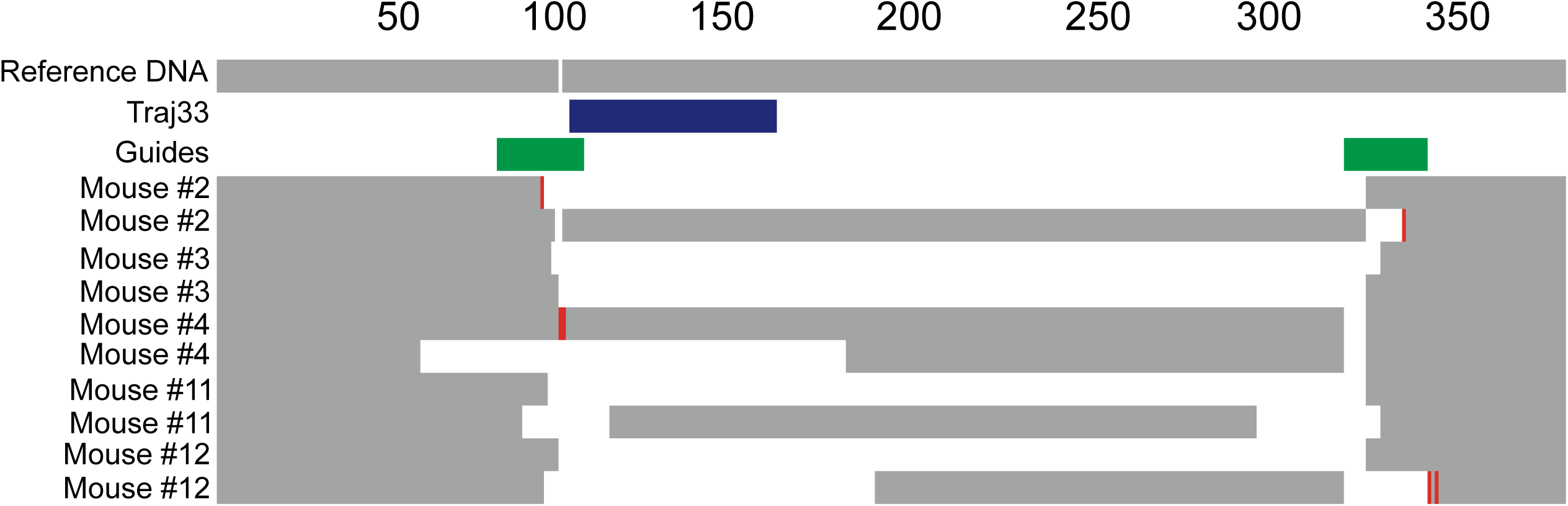
Generation of CC011 Traj33−/− mice. Shown are the top 2 NGS-derived genomic DNA sequences of the 5 CC011-Traj33−/− mice generated by CRISPER/Cas9-mediated knock out of the Traj33 gene. Purple denotes Traj33 nucleotide sequence, green indicates nucleotide sequence of guide RNAs. Grey regions represent consensus sequence aligned to the mm10 reference strain. The white gaps indicate deleted sequence, and red indicates insertions/mutations.

**Figure S9.**
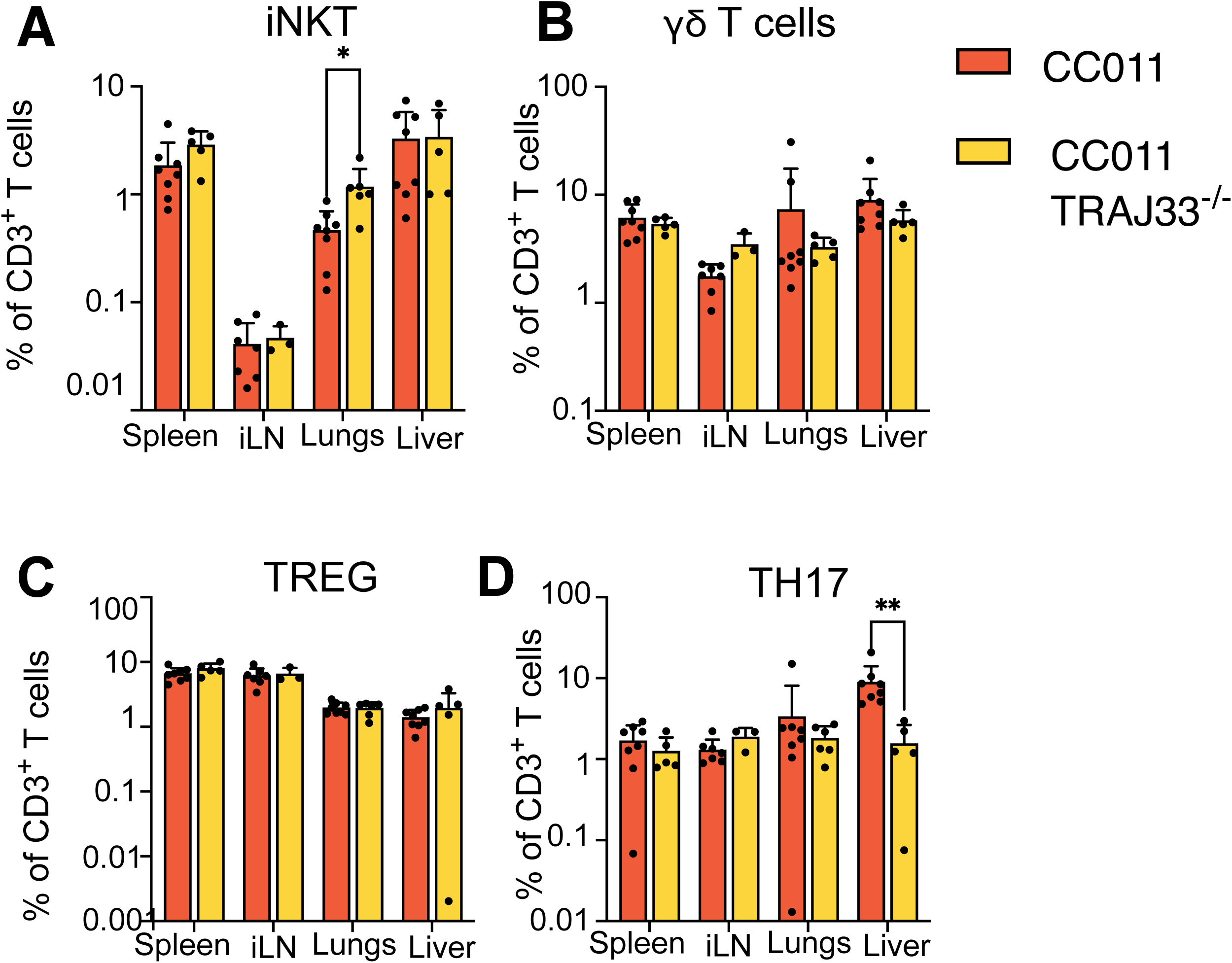
iNKT, γ8 T, T_reg_ and TH17 cell %frequencies in tissues of the MAIT cell deficient CC011-Traj33^−/−^ strain. Mice were aged 25-37 weeks old. *(*P<.05; **P<.01, t-test with Welch correction, n=3-8 mice/group (A-D))*

---

**Table S1:** Thymic MAIT cell frequencies in CC-strains and their founder strains.

| **Strain** | **%MAIT (Mean)** | **SD** | **%MAIT (Mean, SD)** |
| --- | --- | --- | --- |
| CC058/Unc | 0.0005973 | 0.0003481 | 0.0005973,0.0003481 |
| CC033/GeniUnc | 0.000748 | 0.0001886 | 0.000748,0.0001886 |
| CC010/GeniUnc | 0.000827 | 0.0002237 | 0.000827,0.0002237 |
| CC013/GeniUnc | 0.000973 | 0.0005359 | 0.000973,0.0005359 |
| CC029/Unc | 0.001091 | 0.0002667 | 0.001091,0.0002667 |
| CC023/GeniUnc | 0.001091 | 0.0002667 | 0.001091,0.0002667 |
| CC045/GeniUnc | 0.001357 | 0.0008179 | 0.001357,0.0008179 |
| CC021/Unc | 0.00137 | 0.0004258 | 0.00137,0.0004258 |
| 129S1/SvlmJ | 0.001397 | 0.0003296 | 0.001397,0.0003296 |
| CC060/Unc | 0.001416 | 0.0007099 | 0.001416,0.0007099 |
| CC084/TauUnc | 0.0015 | 0.0005567 | 0.0015,0.0005567 |
| CC005/TauUnc | 0.001585 | 0.001049 | 0.001585,0.001049 |
| CC036/UncJ | 0.00173 | 0.0004093 | 0.00173,0.0004093 |
| CC078/TauUnc | 0.001733 | 0.0005613 | 0.001733,0.0005613 |
| CC016/GeniUnc | 0.00184 | 0.0004303 | 0.00184,0.0004303 |
| CAST/EiJ | 0.00206 | 0.0007139 | 0.00206,0.0007139 |
| CC071/TauUnc | 0.00212 | 0.000522 | 0.00212,0.000522 |
| CC072/TauUnc | 0.002533 | 0.00087 | 0.002533,0.00087 |
| CC026/GeniUnc | 0.002183 | 0.001209 | 0.002183,0.001209 |
| CC043/GeniUnc | 0.002227 | 0.0006494 | 0.002227,0.0006494 |
| CC075/Unc | 0.00225 | 0.000383 | 0.00225,0.000383 |
| CC057/Unc | 0.002297 | 0.0009942 | 0.002297,0.0009942 |
| C57BL/6J | 0.002363 | 0.0003811 | 0.002363,0.0003811 |
| CC037/TauUnc | 0.00252 | 0.001621 | 0.00252,0.001621 |
| CC007/Unc | 0.00255 | 0.0008354 | 0.00255,0.0008354 |
| PWK/PhJ | 0.002583 | 0.000415 | 0.002583,0.000415 |
| CC079/TauUnc | 0.002817 | 0.001319 | 0.002817,0.001319 |
| NZO/HILtJ | 0.002817 | 0.0008979 | 0.002817,0.0008979 |
| CC012/GeniUnc | 0.002872 | 0.00278 | 0.002872,0.00278 |
| CC039/Unc | 0.002907 | 0.0009868 | 0.002907,0.0009868 |
| CC041/TauUnc | 0.003073 | 0.002095 | 0.003073,0.002095 |
| CC061/GeniUnc | 0.003213 | 0.001255 | 0.003213,0.001255 |
| CC019/TauUnc | 0.00374 | 0.001255 | 0.00374,0.001255 |
| CC024/GeniUnc | 0.003777 | 0.001666 | 0.003777,0.001666 |
| CC053/Unc | 0.003971 | 0.003 | 0.003971,0.003 |
| CC032/GeniUnc | 0.00442 | 0.001308 | 0.00442,0.001308 |
| CC025/GeniUnc | 0.004467 | 0.001163 | 0.004467,0.001163 |
| NOD/ShiLtJ | 0.004748 | 0.002459 | 0.004748,0.002459 |
| CC049/TauUnc | 0.004782 | 0.004802 | 0.004782,0.004802 |
| CC074/Unc | 0.004813 | 0.00174 | 0.004813,0.00174 |
| CC006/TauUnc | 0.005047 | 0.001241 | 0.005047,0.001241 |
| CC009/Unc | 0.00523 | 0.001696 | 0.00523,0.001696 |
| CC001/Unc | 0.00527 | 0.004966 | 0.00527,0.004966 |
| CC083/UncJ | 0.005437 | 0.001613 | 0.005437,0.001613 |
| CC002/Unc | 0.00578 | 0.001118 | 0.00578,0.001118 |
| CC035/Unc | 0.00604 | 0.00108 | 0.00604,0.00108 |
| CC003/Unc | 0.006313 | 0.00205 | 0.006313,0.00205 |
| CC028/GeniUnc | 0.00636 | 0.003472 | 0.00636,0.003472 |
| A/J | 0.008337 | 0.001445 | 0.008337,0.001445 |
| CC017/UncJ | 0.00926 | 0.002394 | 0.00926,0.002394 |
| CC040/TauUnc | 0.00939 | 0.001455 | 0.00939,0.001455 |
| CC018/UncJ | 0.01116 | 0.005359 | 0.01116,0.005359 |
| CC004/TauUnc | 0.01223 | 0.00225 | 0.01223,0.00225 |
| WSB/EiJ | 0.01224 | 0.003952 | 0.01224,0.003952 |
| CC068/TauUnc | 0.01243 | 0.007884 | 0.01243,0.007884 |
| CC059/TauUnc | 0.01517 | 0.003869 | 0.01517,0.003869 |
| CC030/GeniUnc | 0.04733 | 0.01286 | 0.04733,0.01286 |
| CC011/Unc | 0.07667 | 0.02344 | 0.07667,0.02344 |

**Table S2:**
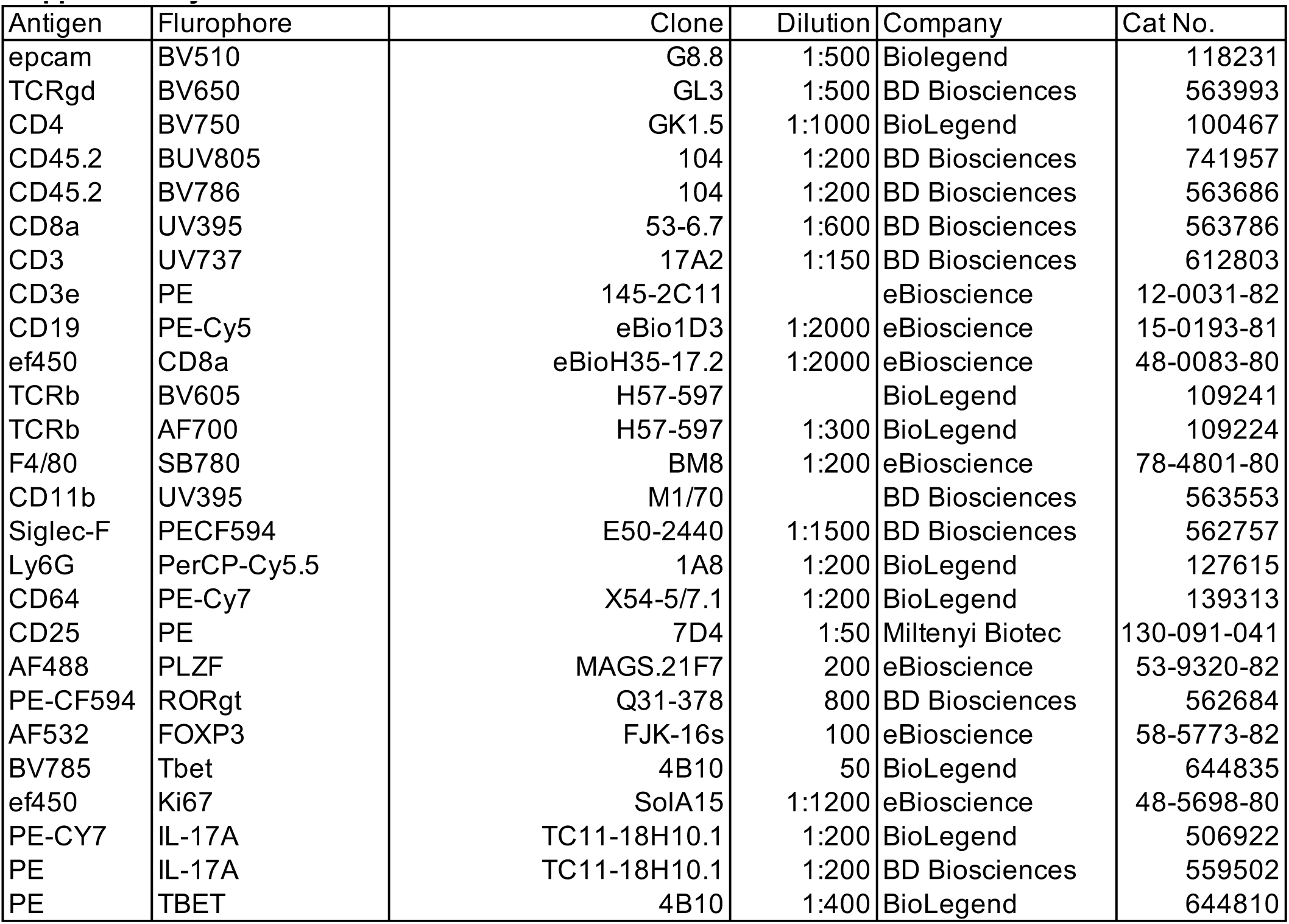
Antibodies and tetramer reagents for surface phenotype and intracellular staining.

| Antigen | Fluorophore | Clone | Dilution | Company | Cat No. |
| --- | --- | --- | --- | --- | --- |
| epcam | BV510 | G8.8 | 1:500 | Biolegend | 118231 |
| TCRgd | BV650 | GL3 | 1:500 | BD Biosciences | 563993 |
| CD4 | BV750 | GK1.5 | 1:1000 | BioLegend | 100467 |
| CD45.2 | BUV805 | 104 | 1:200 | BD Biosciences | 741957 |
| CD45.2 | BV786 | 104 | 1:200 | BD Biosciences | 563686 |
| CD8a | UV395 | 53-6.7 | 1:600 | BD Biosciences | 563786 |
| CD3 | UV737 | 17A2 | 1:150 | BD Biosciences | 612803 |
| CD3e | PE | 145-2C11 |  | eBioscience | 12-0031-82 |
| CD19 | PE-Cy5 | eBio1D3 | 1:2000 | eBioscience | 15-0193-81 |
| ef450 | CD8a | eBioH35-17.2 | 1:2000 | eBioscience | 48-0083-80 |
| TCRb | BV605 | H57-597 |  | BioLegend | 109241 |
| TCRb | AF700 | H57-597 | 1:300 | BioLegend | 109224 |
| F4/80 | SB780 | BM8 | 1:200 | eBioscience | 78-4801-80 |
| CD11b | UV395 | M1/70 |  | BD Biosciences | 563553 |
| Siglec-F | PECF594 | E50-2440 | 1:1500 | BD Biosciences | 562757 |
| Ly6G | PerCP-Cy5.5 | 1A8 | 1:200 | BioLegend | 127615 |
| CD64 | PE-Cy7 | X54-5/7.1 | 1:200 | BioLegend | 139313 |
| CD25 | PE | 7D4 | 1:50 | Miltenyi Biotec | 130-091-041 |
| AF488 | PLZF | MAGS.21F7 | 200 | eBioscience | 53-9320-82 |
| PE-CF594 | RORgt | Q31-378 | 800 | BD Biosciences | 562684 |
| AF532 | FOXP3 | FJK-16s | 100 | eBioscience | 58-5773-82 |
| BV785 | Tbet | 4B10 | 50 | BioLegend | 644835 |
| ef450 | Ki67 | SolA15 | 1:1200 | eBioscience | 48-5698-80 |
| PE-CY7 | IL-17A | TC11-18H10.1 | 1:200 | BioLegend | 506922 |
| PE | IL-17A | TC11-18H10.1 | 1:200 | BD Biosciences | 559502 |
| PE | TBET | 4B10 | 1:400 | BioLegend | 644810 |

**Table S3:**
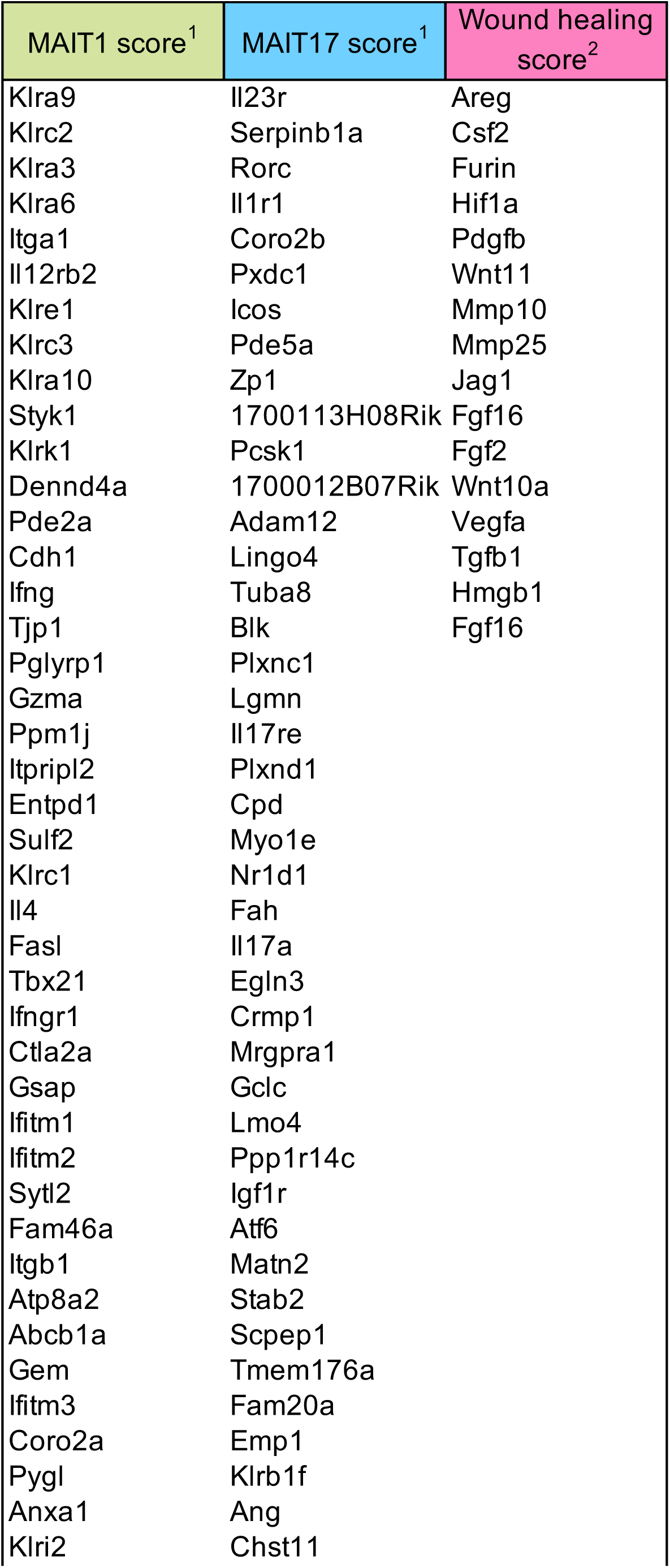
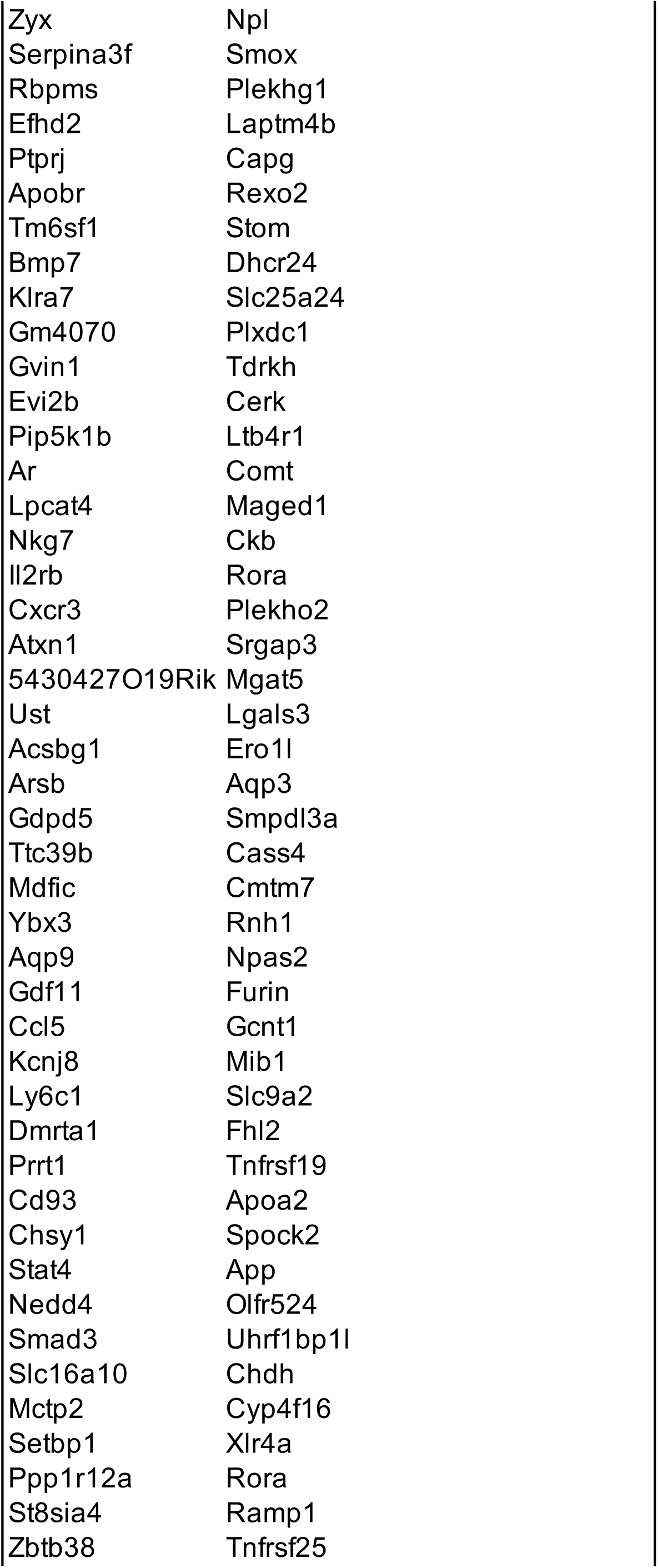
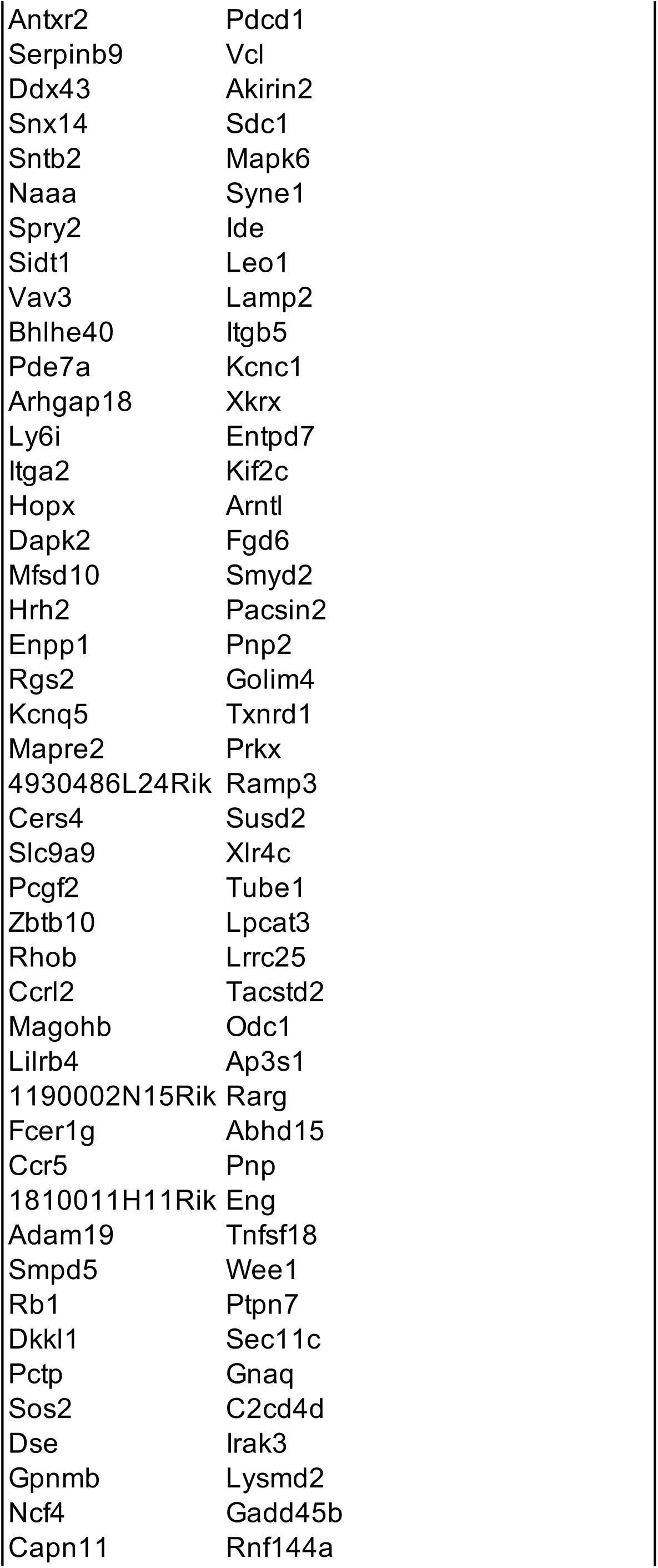
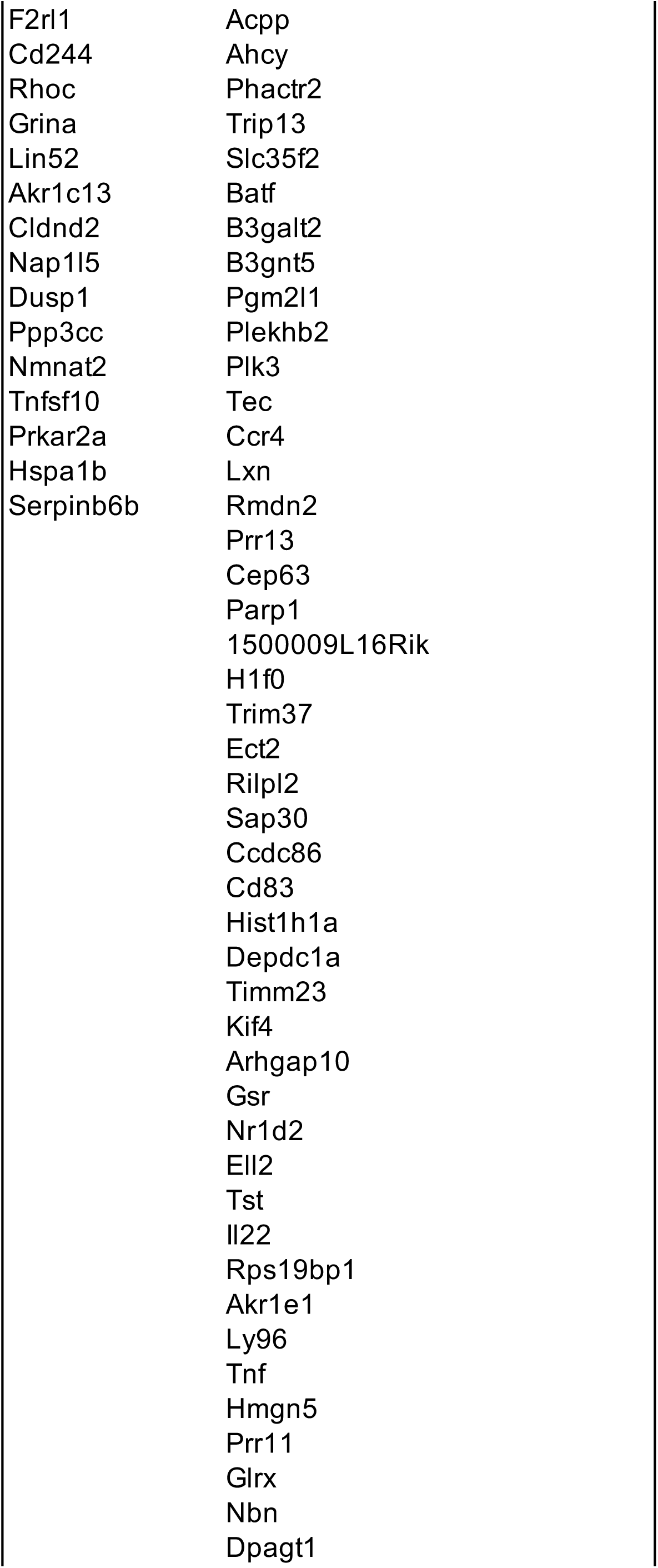
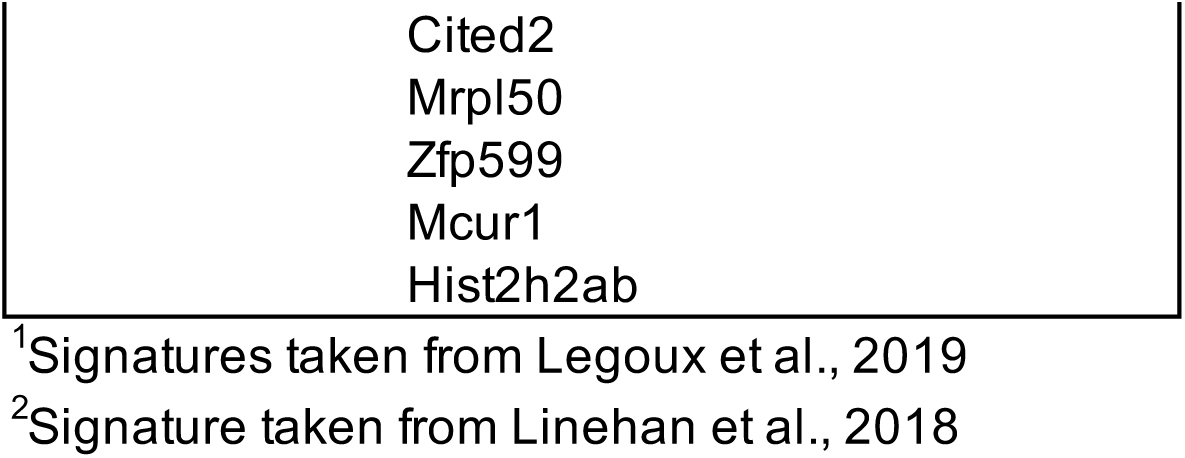
Colon gene signature of MAIT cells (Differentially expressed genes between colon and spleen, Seurat function FindAllMarkers)

**Table S4:**
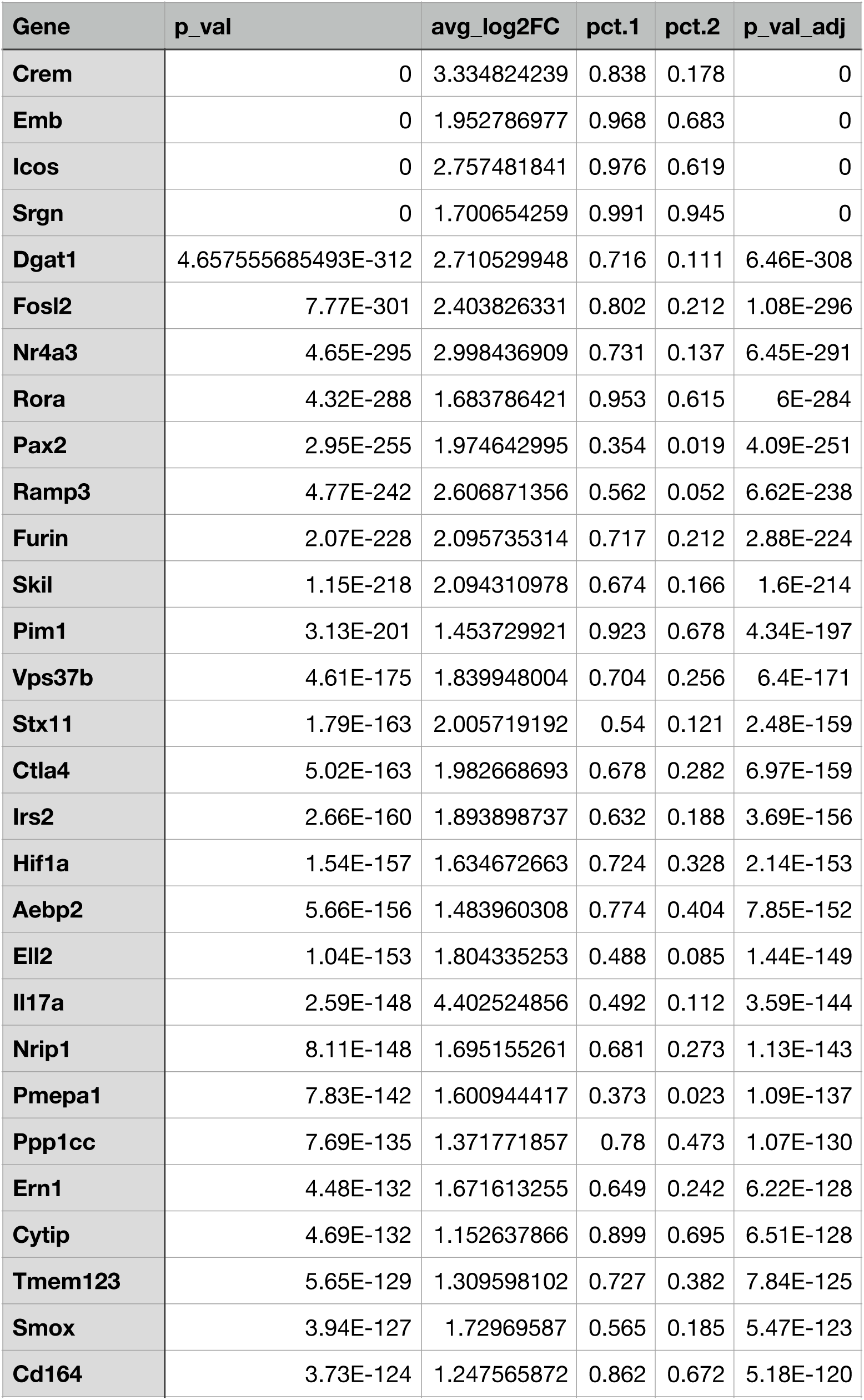
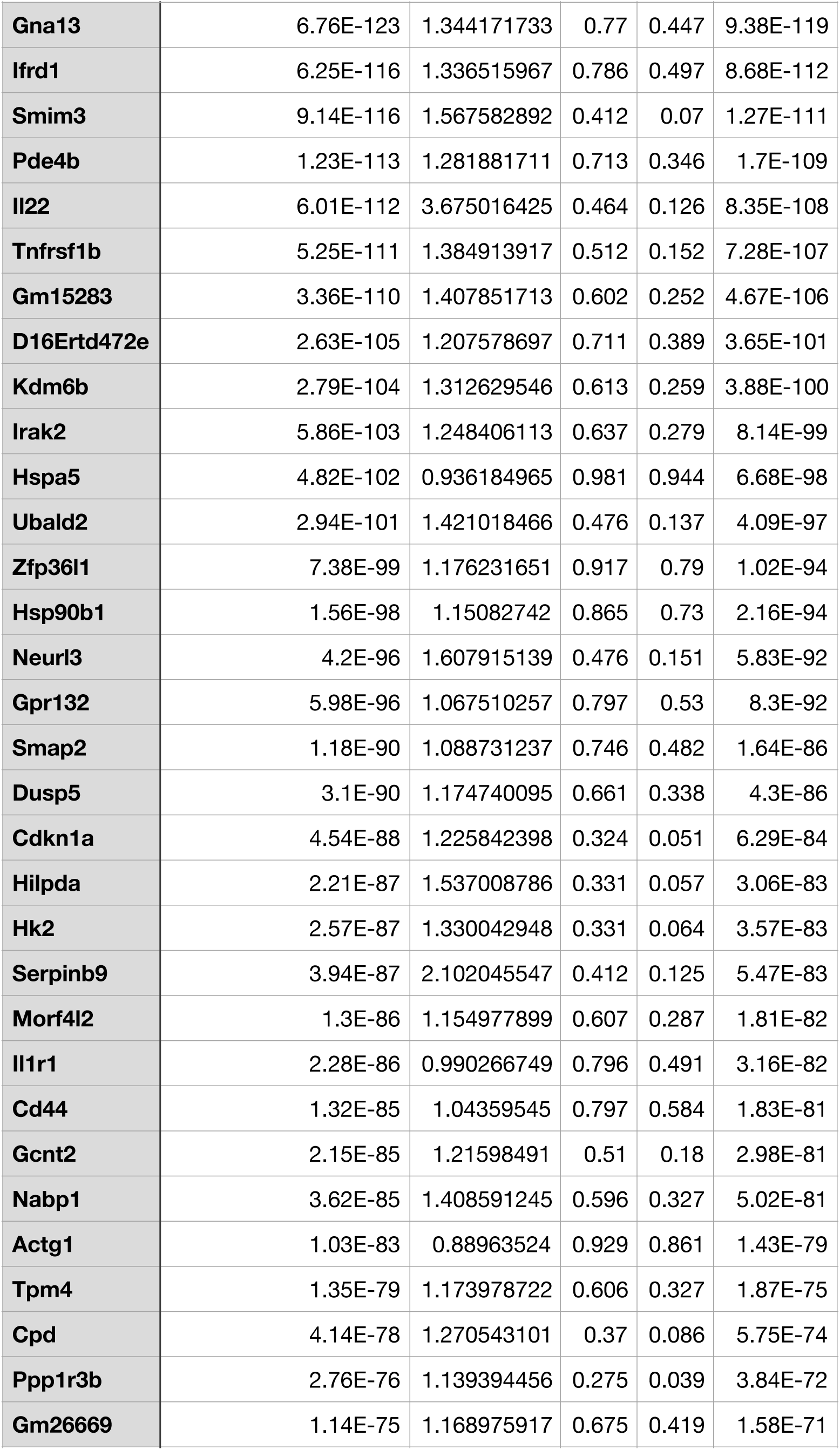
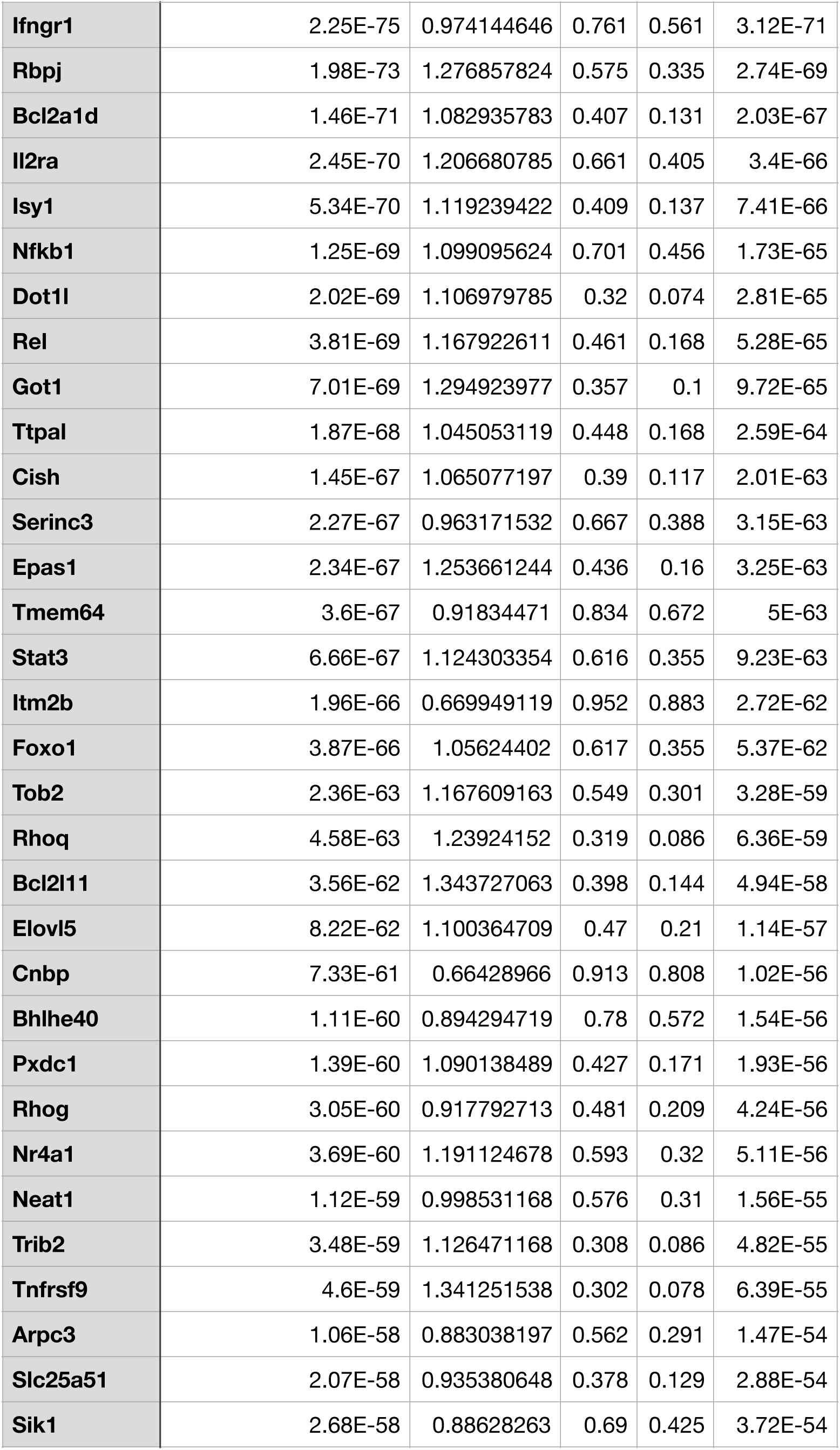
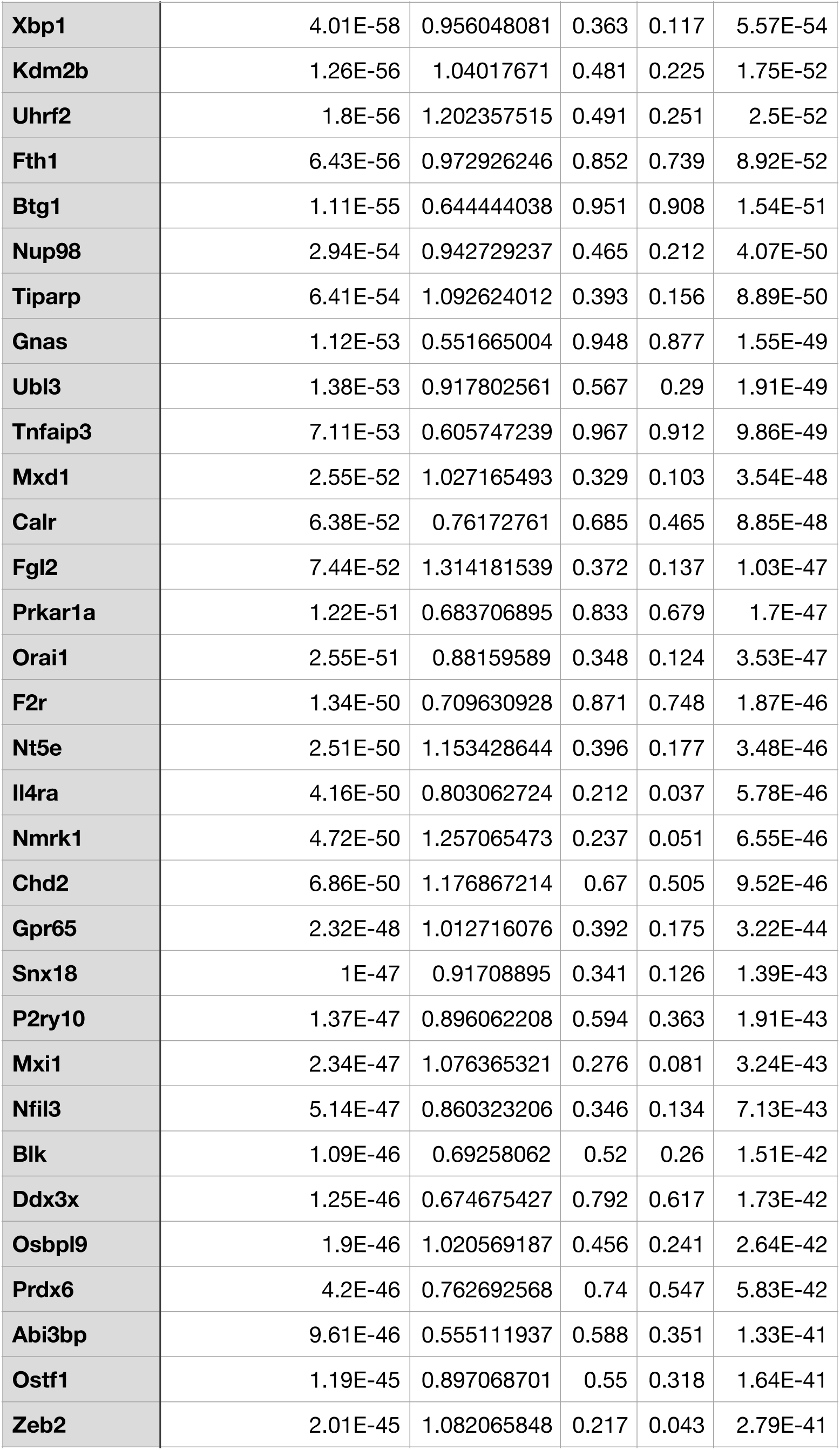
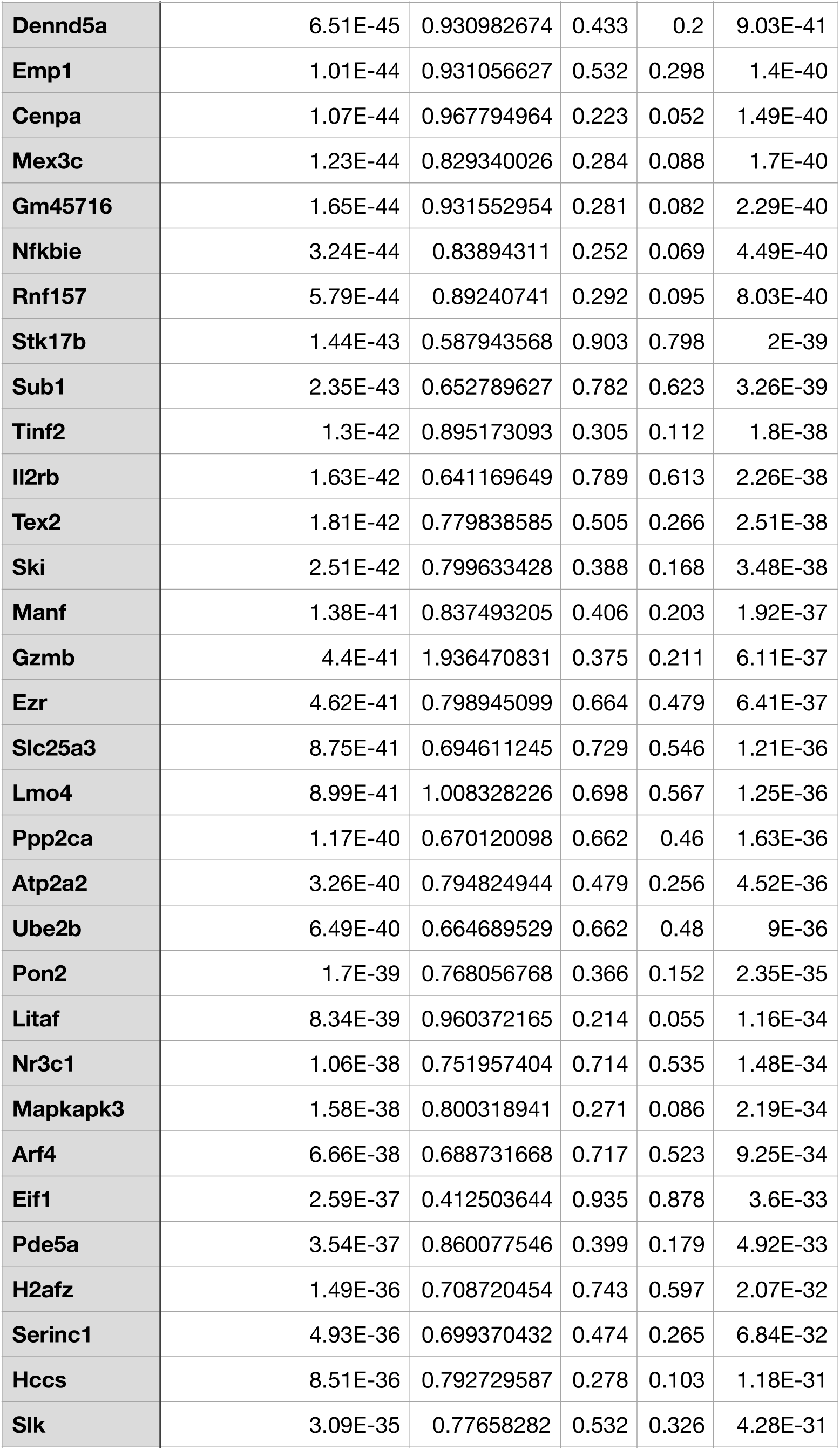
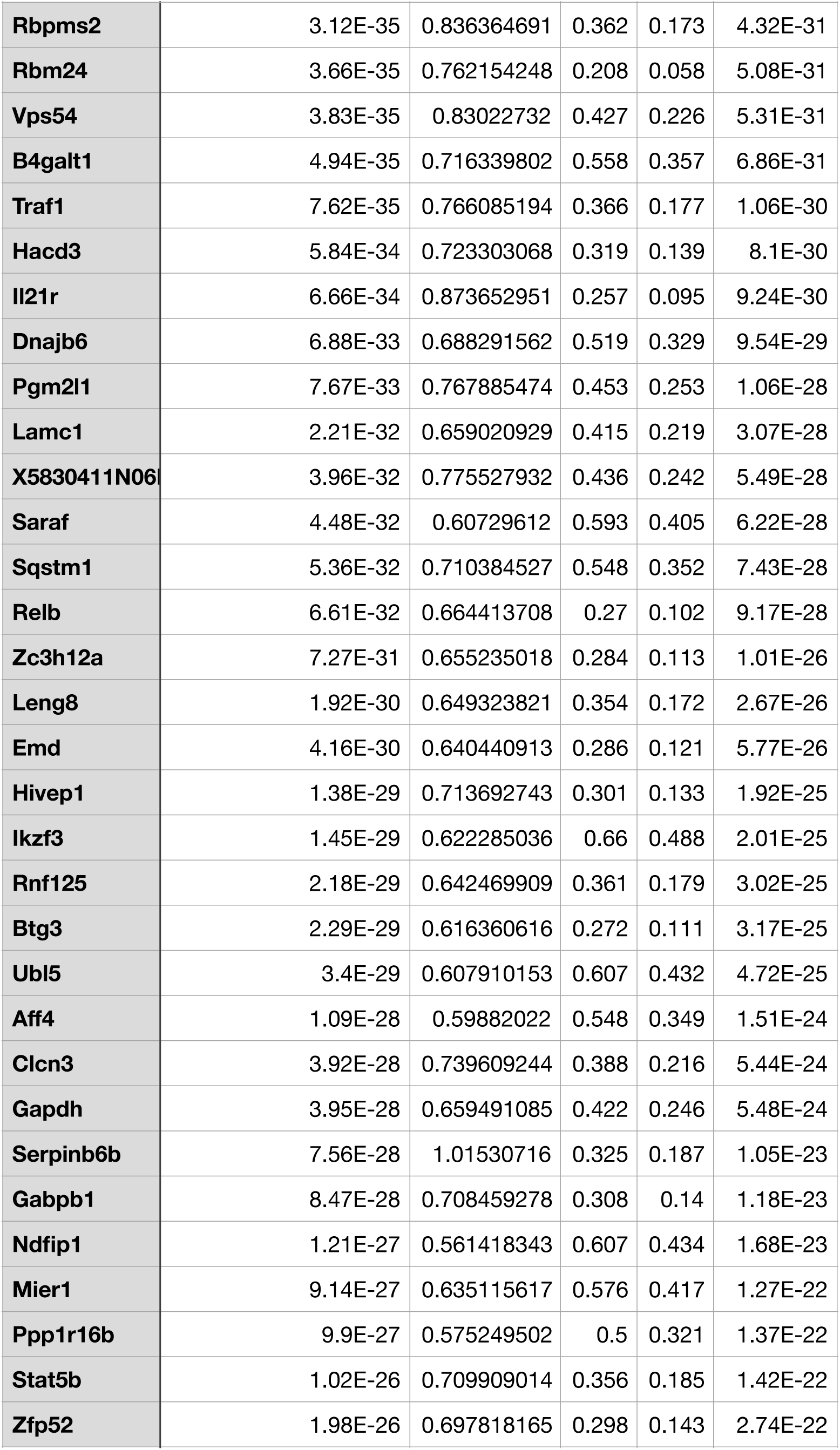
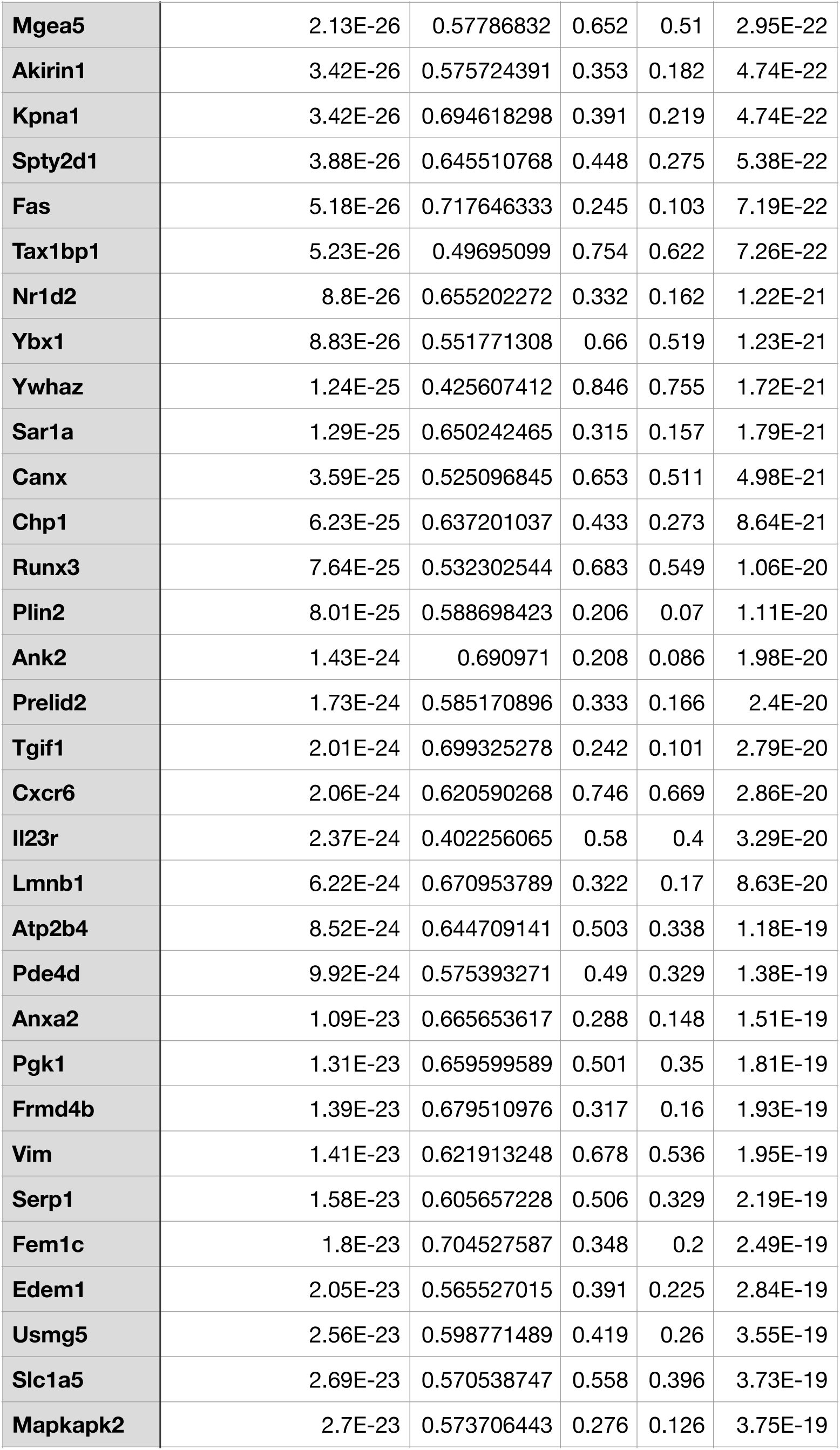
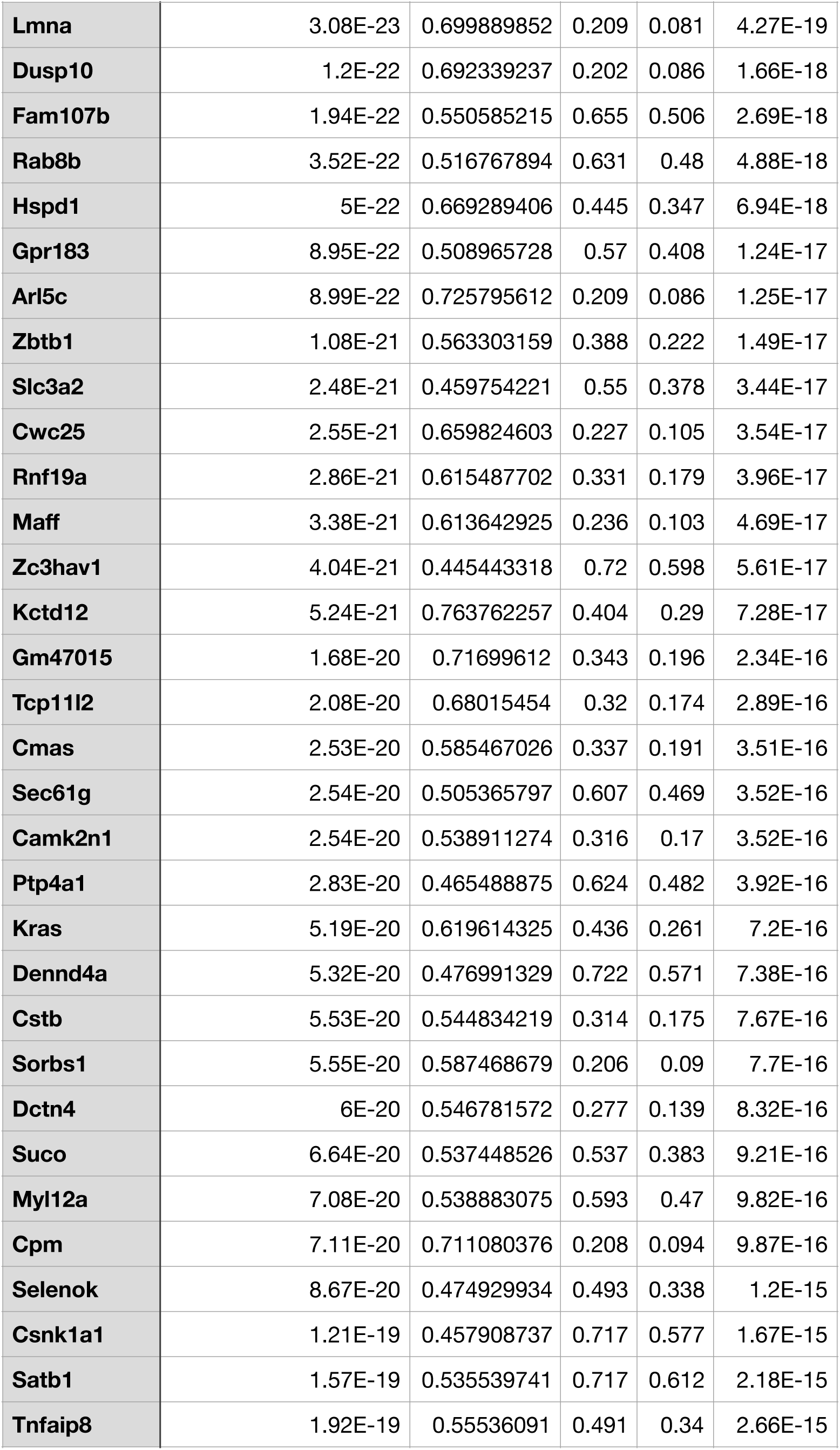
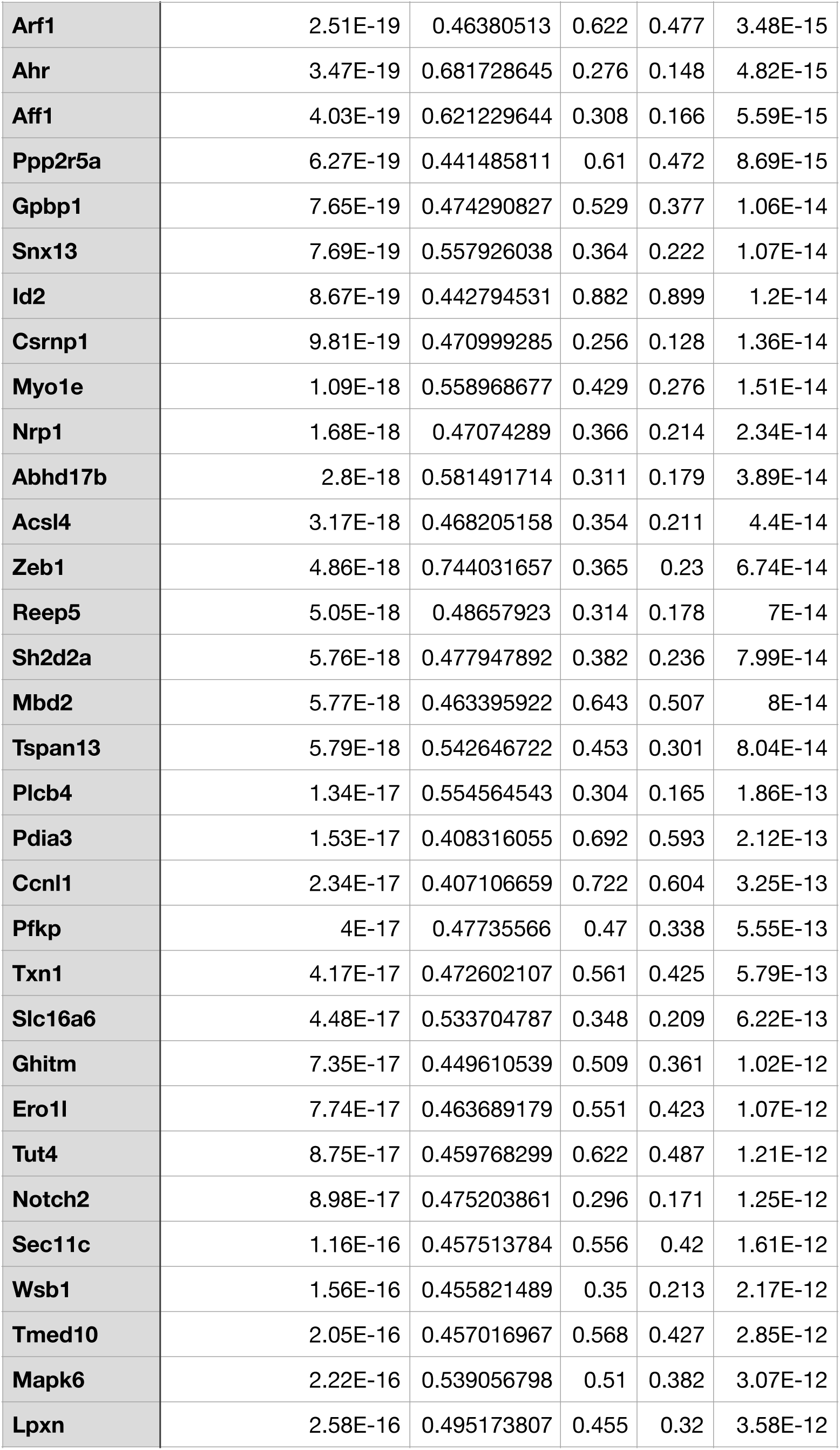
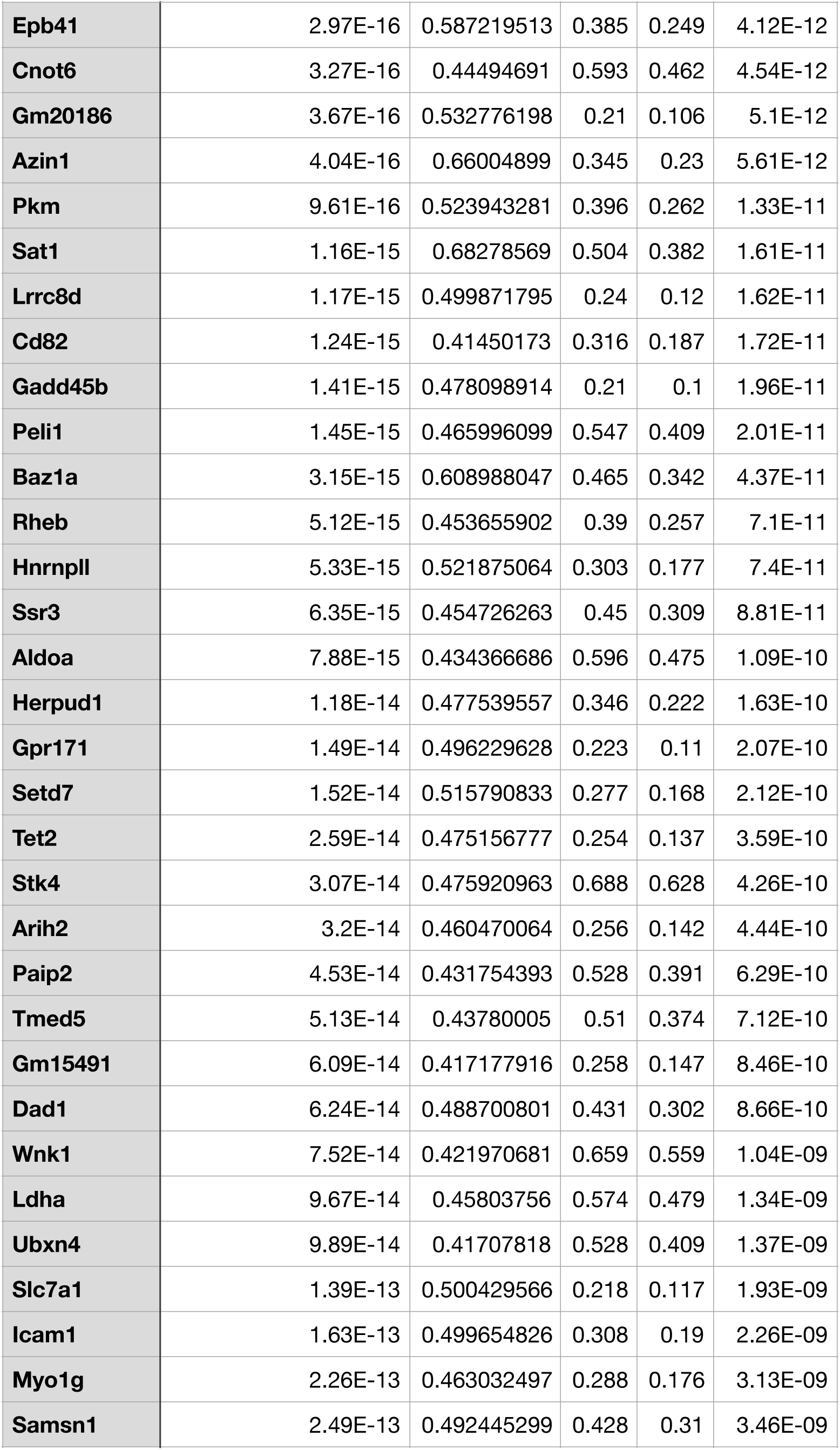
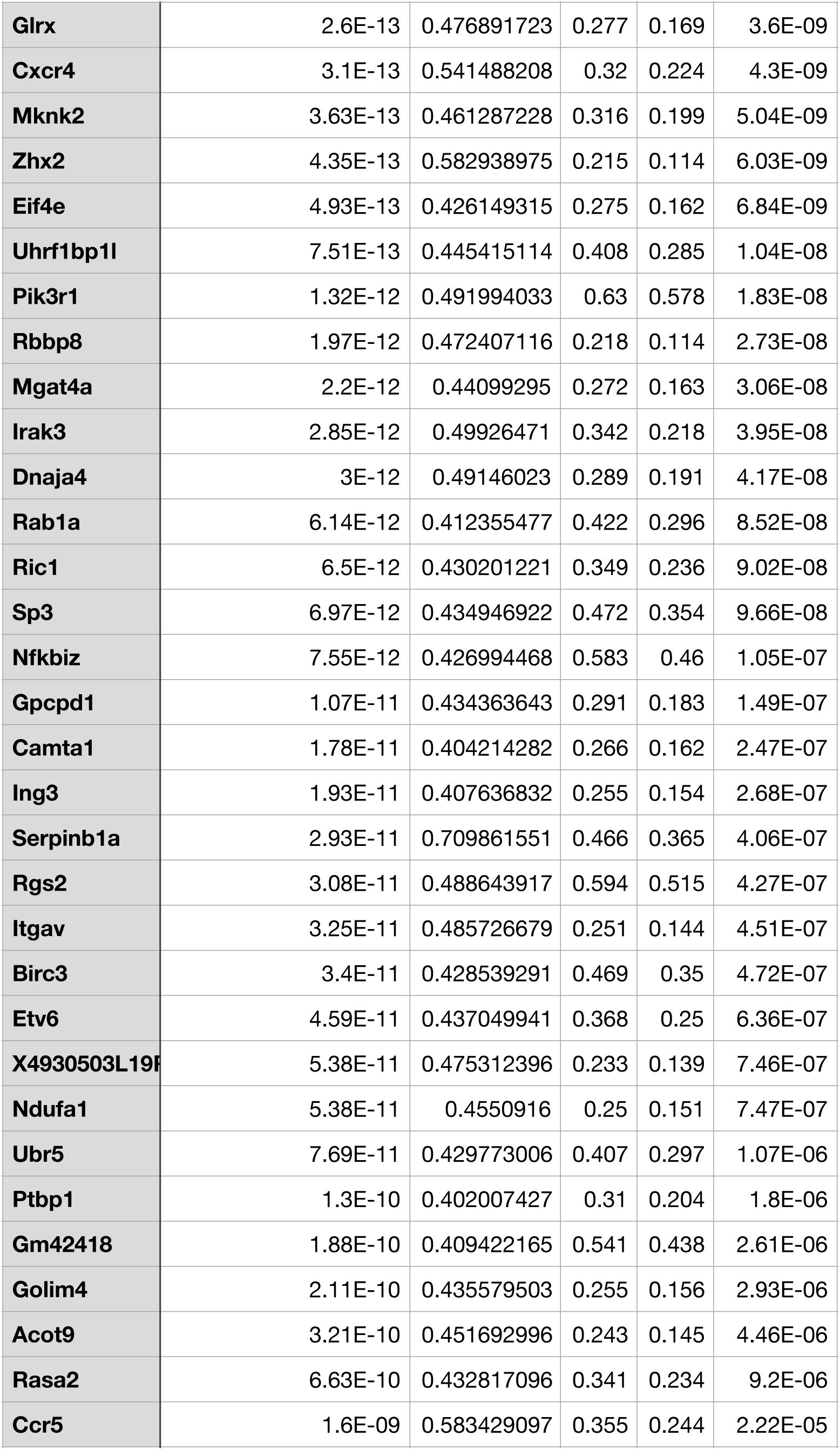
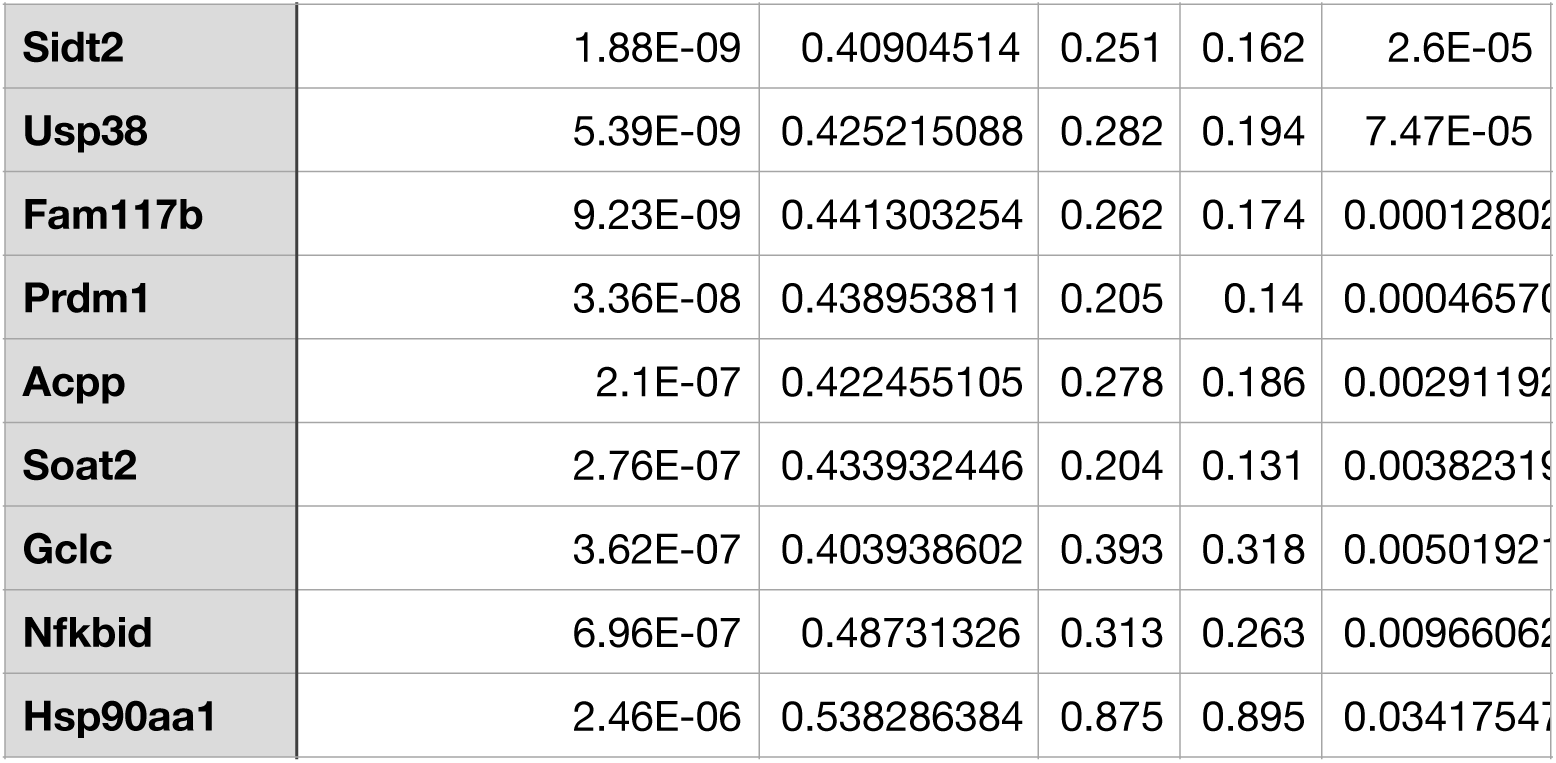
colon_signature.

## Notes

### Competing Interest Statement

The authors have declared no competing interest.

